# Dissecting *ARL15* Function in Rheumatoid Arthritis: Insights from *Ex Vivo* and *In Vitro* Synovial Fibroblast Models

**DOI:** 10.1101/2025.06.30.662131

**Authors:** Sujit Kashyap, Anuj Kumar Pandey, Paritosh Kumar, Maumita Kanjilal, Uma Kumar, Thelma BK

**Affiliations:** Department of Genetics University of Delhi South Campus, New Delhi, India; Rheumatology Department, All India institute of Medical Sciences, New Delhi, India; Feinberg School of Medicine, Northwestern University, Chicago, USA

## Abstract

*ARL15*, coding for a small GTPase was identified as a non-HLA susceptibility gene in rheumatoid arthritis (RA) through a GWAS in a North Indian cohort. Serum adiponectin and ARL15 levels were higher in RA patients with the associated genotype. The present study aimed to delineate the functional role of *ARL15* in RA pathobiology using gene knockdown (KD) combined with transcriptomic profiling in both *ex-vivo* RA synovial fibroblasts (RASF) and *in vitro* MH7A cell lines. In RASF, *ARL15* KD led to the downregulation of *COMP*-an extracellular matrix stabilizer linked to severe RA-alongside upregulation of adiponectin and IFN response genes such as *IFI6* and *USP18*. Furthermore, upregulation of *NPTX1* and *MX1*, previously associated with disease modulation and treatment response was observed. Downregulation of *CTGF, CD248*, and *PTX3* suggested involvement of *ARL15* in inflammation and RA-associated cardiovascular risk. In contrast, *ARL15* KD in MH7A cells displayed distinct gene signatures with upregulated cytokines (*IL1A, IL8, CXCLs*) and downregulated inflammatory regulators (*DOCK2, TLR4, TGFB2*), reflecting an inflammatory bias distinct from the patient-derived RASF. This divergence highlights the limitations of immortalized cell models in capturing patient heterogeneity and disease complexity. However, the dual-system approach underscores the multifaceted role *of ARL15* in regulating connective tissue architecture, inflammation, and immune response. These key findings position *ARL15* as a promising therapeutic target, warranting further investigation in RA animal models and genomic medicine. Taken together, this work provides a compelling rationale to pursue ARL*15* targeted interventions in RA management.

## Introduction

Rheumatoid arthritis (RA) is a common, chronic systemic autoimmune disease characterized by nonspecific inflammation of the peripheral joints, which leads to synovial damage, pain, stiffness, and reduced functional capacity (1, 2). Synovial joints are specialized organs designed to meet unique biomechanical and physiological demands at specific anatomical locations. These joints comprise structural elements such as articular cartilage, synovial membrane (a delicate structure consisting of one or two cell layers of synovial fibroblasts and tissue-resident macrophages), ligaments and menisci, which collectively support joint function. The synovium contains a structurally distinct lining layer that interfaces with the synovial fluid(3). A hallmark feature of RA pathology is the marked hyperplasia of this synovial lining due to autoimmune activation. Synovial fibroblasts (SF), commonly referred to as rheumatoid arthritis synovial fibroblasts (RASF) in RA patients, are among the most destructive cell types in the synovium. These cells exhibit resistance to apoptosis, leading to an increase in their numbers and contributing to synovial hyperplasia, a critical factor in disease progression (4).

RA manifests significant clinical heterogeneity, complicating diagnosis, prognosis and treatment. Progression of RA varies with patients as few achieve remissions or low disease activity whereas others develop severe joint deformity and functional disability (5). The etiology of RA involves a complex interplay of genetic and non-genetic factors. Non-genetic factors implicated range from smoking, air pollution, obesity to alcohol consumption etc. (6, 7) but mechanism of their contribution or pathways adopted remain largely unexplained. Conversely, over 100 susceptibility genes have been identified based on meta-analyses of GWAS data in populations of European, Asian, and other ancestries (8-10). These findings clearly indicate genetic heterogeneity and likely population specific susceptibility in RA.

In the only GWAS in RA from India to date, we identified *ARL15*, a non-HLA gene associated with the disease (11). Preliminary characterization of *ARL15*, a member of the ARF family of small GTPases using RASF and a knock down (KD) approach showed a significant decrease in the invasion and migration properties of RASF. It was also demonstrated that *ARL15* action is independent of (12) (13). In animal models of RA, the inhibition of ARF proteins alleviated inflammation and disease severity (14). *ARL15* has been associated with the age of onset for alcohol dependence (15), which, in turn, correlates with RA risk (16) and RA is closely linked to cardiovascular disease (CVD) risk factors. *ARL15* is also associated with metabolic traits, including HDL cholesterol concentration, insulin resistance, coronary heart disease, fasting insulin, and triglyceride levels (17). However, the precise mechanisms and pathways underlying the role of this novel susceptibility gene in RA and other complex traits remain unclear.

In this study, we sought to gain deeper insights into the role of *ARL15* in RA biology by employing *ARL15* KD and hypothesis-free transcriptomics strategy using both patient derived RASF cells and MH7A, a RASF derived cell line. Analyses revealed i) differential expression of functionally relevant genes ii) differences in these between the two experimental RASF samples; iii) similarities and differences in findings between RASF samples and MH7A cells; and iv) most importantly, novel insights into the likely role of *ARL15* in RA.

## Materials and methods

### Recruitment of study subjects and collection of target tissues

This study was carried with institutional ethical committee clearance from both participating centers. Patients with RA were diagnosed at Rheumatology unit All India Institute of Medical Sciences, New Delhi (by UK) in accordance with the American college of rheumatology/ EULAR criteria (18). Synovial fluid/tissue samples from study subjects undergoing total knee replacement surgery at the orthopedic surgery unit at the same hospital were collected with informed consent. Samples from osteoarthritis (OA) patients recruited the same way were used as controls.

### Generation of patient derived synovial fibroblasts (RASF)

Tissue samples from five RA patients (synovial fluid from patient id # RA1, RA3 and RA6; synovial tissue from id # RA4 and RA6) and two OA patients (synovial fluid from id # OA1 and synovial tissue from id # OA2) were obtained during total knee replacement surgery. Synovial fibroblasts from these two tissues were cultured as described elsewhere (19) Briefly, synovial fluid was aspirated and transferred to a heparin vacutainer and immediately transported on ice to the laboratory. The sample was centrifuged at 1100 rpm for 10 min and the pellet was seeded in a T-25 flask and grown at 37^°^C, in a 5% CO_2_incubator in complete medium (with 15% fetal bovine serum, 1% non-essential amino acids and 1% penicillin-streptomycin solution). Synovial tissue collected was transported on ice to the lab and first minced into small pieces and then incubated with collagenase (C2139, SIGMA) in DMEM (1199504, Thermofisher, USA) overnight at 37^°^ C. The suspension was then centrifuged at 1200 rpm for 15 and pellet was dissolved in complete medium (DMEM, 10% fbs, 1% penicillin-streptomycin solution) and seeded in a T-25 flask. Cells were then grown at 37^°^ C and sub-passaging for both tissue and fluid derived synovial fibroblasts was done as required and aliquots were also frozen for future use. Cultures with ∼80% confluence after third passage were used for all the experiments as previously described (13).

### Immortalized synovial fibroblast (MH7A cell line) culture

MH7A synovial fibroblast cell line (RCB1512) was procured from RIKEN cell bank, Japan and used for *ARL15* KD experiments. Briefly, cells were cultured in RPMI 1640 media (11875093, Thermo Fisher Scientific, USA) supplemented with 10% heat-inactivated Fetal Bovine Serum (10100147, Thermo Fisher Scientific, USA), 1% penicillin-streptomycin (10,000 U/mL) (15140122, Thermo Fisher Scientific, USA), and were incubated at 37^°^C with 5% CO2 for maintenance and subsequent experiments.

### Knockdown of ARL15 using siRNA

siRNA for *ARL15* and scrambled siRNAs to be used as negative control (NC) were the same as described previously (Kashyap et al., 2018). RASF and OASF samples were seeded at a density of 1.5x10^6^ in six well plates. 24 hours prior to transfection, cells were incubated in antibiotic and serum free medium. 10nM of *ARL15* siRNA (A50284F6, Thermo Fisher Scientific, USA) and scrambled siRNA as NC (AM4611, Thermo Fisher Scientific, USA) were used for respective transfections using lipofectamine 2000 (11668019, Thermo Fisher Scientific, USA). Six hours later the medium containing siRNA was replaced by normal growth medium, and cells were grown for 24 hrs. at 37^°^C and harvested for RNA isolation. RASF cultures with ∼80% confluence treated with only lipofectamine, and Opti-MEM culture medium were considered as wild type controls. MH7A cells were grown in a T-25 flask to ∼ 90-95% confluency. Adherent cells were dislodged using 0.25 % trypsin-EDTA solution (25200072, Thermo Fisher Scientific, USA) and resuspended in 1000 μL of reduced serum medium (Opti-MEM media, 31985070, Thermo Fisher Scientific, USA). ∼10 μL of resuspended cells stained with trypan blue (T10282, Thermo Fisher Scientific, USA) were used to count the total number of live/dead cells/mL using an automated cell counter (AMQAF1000, Thermo Fisher Scientific, USA). Around 80-100 μL (∼0.2-0.3x10^5^ cells/mL) of resuspended cells per well of a six-well plate were used for reverse transfection. Like for RASF, 10nM of *ARL15* siRNA and scrambled siRNA as NC were used. siRNA-Liposome mixture was prepared accordingly on a per well basis of a six-well plate using Lipofectamine 3000 and P3000 reagents (5 μL each) (L3000015, Thermo Fisher Scientific, USA) and used for KD experiments. For a good transfection efficiency, reverse transfection was performed as per the standard protocol by adding siRNA and Lipofectamine together followed by addition of resuspended cells to the mix. Briefly, Opti-MEM-Lipofectamine mix was added with the siRNA-P3000-Opti-MEM mix in equal (1:1) proportion and incubated for 5 minutes at room temperature. ∼80-100μL of resuspended cells was added to above mix, and the mixture was added to the respective wells having Opti-MEM media up to 2000 μL and incubated for 6 hours at 37^°^ C with 5% CO_2_ for surface attachment, followed by replacement of Opti-MEM media with complete media for further growth.

### Total RNA extraction

Total RNA was extracted from wild-type (WT) and knockdown (KD) RASF/RAST cells using the standard Trizol method (TRI reagent, T9424, Sigma, USA). Briefly, cells were washed with PBS before treatment with Trizol for 5 minutes. RNA was isolated using chloroform extraction, followed by precipitation with isopropanol and a 75% ethanol wash. The final RNA pellet was dissolved in nuclease-free water, and its quantity and quality were assessed using a Nanodrop (Thermo, USA) and a Bioanalyzer (Agilent, USA).

For MH7A cells, RNA isolation was performed using Trizol reagent as above but isolation was done with RNA isolation kit (R2070, Zymo Research, USA). Extracted RNA was quantified using a fluorometry-based method with an RNA quantification kit (Q32852, Thermo Fisher Scientific, USA). RNA samples thus isolated were used for transcriptome sequencing.

### Transcriptome sequencing

RASF RNA isolated from RA3 (synovial fluid), RA6 (synovial tissue), and OASF from OA1 (synovial fluid) cells, assessed for quality and integrity (RIN ≥ 8), were used for transcriptome sequencing. Libraries were prepared using 400 ng of RNA per sample with the TruSeq RNA Library Prep Kit (Illumina, USA), following the manufacturer’s protocol and sequencing performed in paired-end mode (2×101 bp) on an Illumina HiSeq 2000, using a commercial facility (MedGenome, Bangalore, India).

For the MH7A cell line, RNA isolated from two independent samples from passages (P7 and P10) following confirmation of knockdown were used for transcriptome sequencing. All QC-passed samples with RIN ≥ 7.2 were used for library preparation. Library preparation was performed using the Ribo-Zero rRNA removal kit (Illumina, Inc., USA) as per the manufacturer’s protocol and sequencing was done as above.

### Transcriptome data analysis

The raw sequencing data in FASTQ format were checked for quality using FastQC. Adapter removal and trimming of low-quality bases were performed using Cutadapt. For differential gene expression analysis in RASF samples, high-quality reads were aligned to the human reference genome (hg19, UCSC Genome Browser) using TopHat v2.0.8 (20). The aligned reads were assembled into transcripts without a reference genome using the Cufflinks software package (v2.0.2) (21). Differential expression between WT and KD samples was determined using the Cuffdiff module in Cufflinks and reported as Fragments Per Kilobase of exon per Million fragments mapped (FPKM). Gene ontology analysis was performed using DAVID Bioinformatics Resources 6.8 (22). Additionally, transcriptome data from two synovial tissue-derived RASF samples—one from a patient and one from a control both of Caucasian origin (23) were included for comparative analyses.

For MH7A cells, raw sequencing data underwent quality assessment using FastQC followed by data filtering, adapter trimming, and contamination removal using Trimmomatic (24). Read alignment, genome mapping, and gene expression quantification were performed using STAR (25). Gene expression levels were represented as FPKM, and differential expression was estimated using DESeq2 (26). Differentially expressed genes with an adjusted p-value (≤ 0.05) and a fold-change of ±1 were considered significant and taken forward for functional annotation. Functional annotation of significantly upregulated or downregulated genes was performed using reported literature in PubMed, as well as the UniProt and GeneCards databases (27). Pathway analysis was carried out as described above for RASF.

## Results

### Confirmation of homogeneity of fluid and tissue derived RASF

RASF cultured from both synovial fluid and synovial tissue exhibited >99% specificity for CD90/Thy-1, with ≤ 0.5% contamination from other cell types, confirming test sample homogeneity (Supplementary Figure 1).

### RASF cells

*ARL15* KD showed a significantly lower expression level of ARL15 in all three samples (two RA and one OA) as compared to the wild type. Differential expression analysis identified 217, 340, and 87 genes (P < 0.05) in the three samples, respectively. Only two genes, *NPTX1* and *MX1*, were common between RA and OA samples, whereas 25 genes including *ARL15* were shared between the two RA samples (Figure 1A, Supplementary tables 1-4). *NPTX1* was found to be upregulated in all three samples, though its expression level was very low in OASF. In contrast, *MX1* was upregulated in RASF but downregulated in OASF. Of the 16 genes common to both RA samples, *COMP, CTGF, KCDN3, PTX3, PRELP, CD248, ACAN, CITED2,* and *NGF* were downregulated, whereas *IFI6, LRRC17, SLC45A2, USP18, COL14A1, CENPK,* and *GPR153* were upregulated. Of note, six other genes exhibited differential regulation between the two RA samples-*ESM1, KRT19, SLN,* and *IL7R* were downregulated, and *MCF2L* and *GRIP1* were upregulated in fluid-derived RASF but showed the exact opposite trend in tissue-derived RASF. These are shown in (Supplementary table 1).The expression of the two most differentially expressed genes, *NPTX1* and *COMP* was validated by qPCR. Significant (P < 0.0001) upregulation of *NPTX1* and downregulation of *COMP* were confirmed in both RASF samples (Figure 1B). Gene Ontology analysis revealed enrichment of genes associated with the extracellular matrix, and the PI3K-AKT pathway emerged as a common pathway in both samples (Figure 2 A, B).

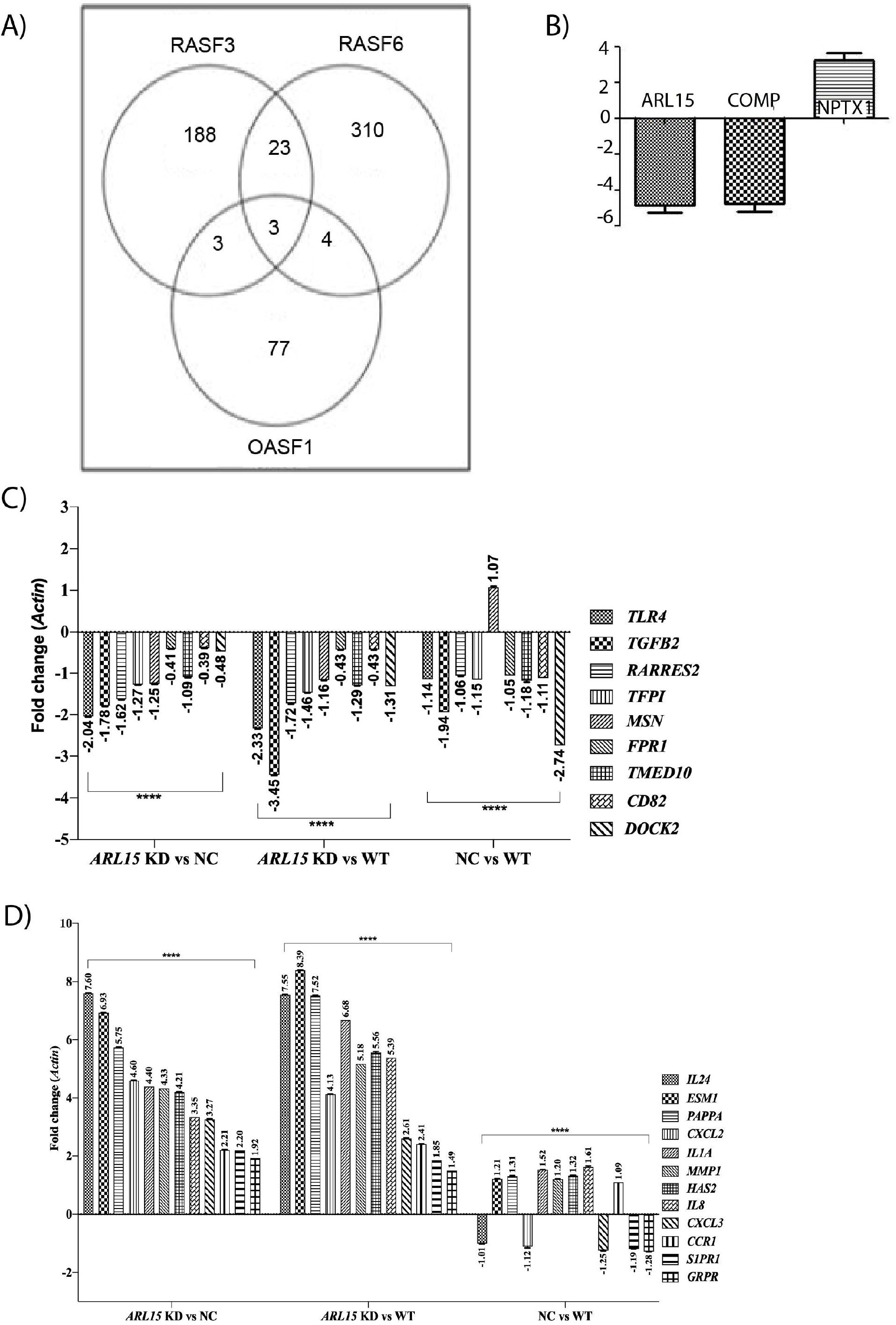

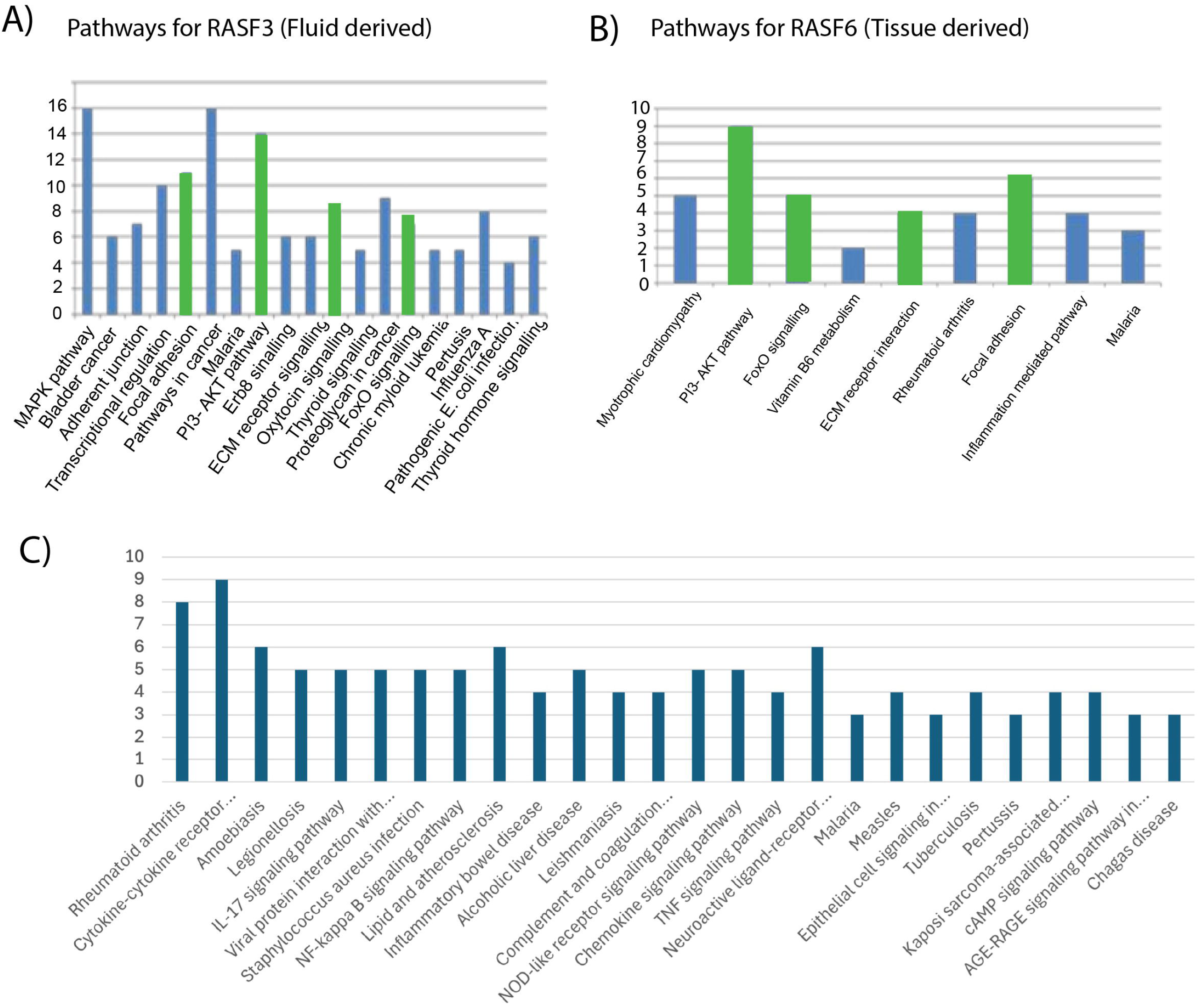

### Increased apoptosis in RASF on *ARL15* KD

The Annexin V staining followed by flow-cytometry in RASF with *ARL15* KD showed ∼10% more cell death (P<0.01) as compared to RASF with WT *ARL15* (Supplementary figure 2).

### MH7A cells

A total of 56 differentially expressed genes (33 upregulated and 23 downregulated) including two long non-coding RNAs identified along with their functional annotation of all are presented (Supplementary table 5). Of these, expression of 21 genes of notable functional relevance including *DOCK2, TGFB2, TLR4, FPR1, TFPI, CD82, RARRES2, MSN,* and *TMED10* (downregulated, Figure 1C); and *CCR1, CXCL2, CXCL3, ESM1, GRPR, HAS2, IL24, IL8, IL1A, MMP1, PAPPA,* and *S1PR1*(upregulated, Figure 1D) was validated through qPCR using the same set of RNA that was used for transcriptome sequencing, with *Actin* and *UBC* as endogenous reference controls (See supplementary information).

### Network analysis of differentially expressed genes

STRING based (Szklarczyk *et al.,* 2019) network analysis was done with 54 differentially expressed genes eliminating PAX8-AS1 and UCA1 (long non-coding RNAs) to identify the gene clusters and the pathways/processes. Two main cytokine-chemokine clusters and keratin protein clusters were observed and *ARL15* is found to interact with *DDIT4L* (based on literature-based text mining). As STRING network analysis did not provide any conclusive idea about the mechanistic/downstream pathway through which *ARL15* functions in the MH7A cell line, DAVID was used for GO and pathway analysis. Again, several inflammation related pathways were observed (Fig 2C). To get more in-depth insight into the role of *ARL15*, all the 56 differentially expressed genes were used for pathway analysis using the IPA tool (Ingenuity Systems, Redwood City, CA; www.qiagen.com/ingenuity). Results based on involvement of 56 genes showed disease and disorders, molecular and cellular functions, and physiological system development and function among the top canonical pathways.

## Discussion

*ARL15* (rs255758 intronic, G>A) was identified as a non-HLA gene associated with RA in the first GWAS that was conducted on a north Indian cohort. Furthermore, in RA patients with the homozygous variant (AA), altered serum adiponectin level was observed (11). Subsequently lower levels of ARL15 in patients with the associated variant genotypes (GA+AA) were documented (28). With hypothesis testing efforts to characterize *ARL15* in RASF using the KD approach, we demonstrated downregulation of *IL6* and upregulation of adiponectin, adiponectin receptor I and TNF independent expression of *ARL15* ; and a reduction in invasion and migration potential of RASF (13, 29). However, the mechanism of action of *ARL15* in RA biology *per se* remains to be known. Therefore,to identify the genes/pathways altered by this small GTPase with likely implications for disease biology, we employed a gene KD approach combined with the hypothesis free transcriptome sequencing in *ex vivo* (patient derived) RASF-both synovial tissue and fluid derived and *in vitro* (MH7A, a well-established cell line) samples. Patient derived RASF serves as a gold standard for studying disease pathology. RASF retains the genetic makeup and primary disease characteristics thus providing a disease comparable physiological milieu. Being derived from different patients, they have the advantage of representing the disease heterogeneity and variability. At the same time, such individual variability and limited life span pose limitations in the use of RASF. It also poses logistical challenges in obtaining and standardizing samples along with ethical and consent-related constraints. In contrast to RASF, immortalized synovial fibroblast (iSF) based cell lines such as MH7A have the advantage of unlimited proliferation, which helps in reproducibility and large-scale experiments; and are easier to manipulate genetically or experimentally as compared to RASF. However, iSF encounter a few limitations such as the potential loss of primary disease characteristics or differentiation state, and importantly, it cannot capture patient-specific genetic or disease-specific clinical heterogeneity. With this rationale and to capture ARL15 biology to the extent possible, we characterized *ARL15* through a knockdown approach followed by transcriptome analysis using both *ex vivo* (RASF) and *in vitro* (MH7A) systems, Insights from genes differentially expressed in these two relevant yet distinctly different experimental systems are discussed.

### RASF findings

Transcriptome data (Supplementary table 1) from the KD experiments in both the RASF samples (RASF3, RASF6) confirmed our previous findings of significant downregulation of *IL6* in synovial fluid derived RASF and a similar trend in tissue derived fibroblast. Adiponectin was not detected in the WT controls but was identified in *ARL15* KD RASF from synovial tissue as mentioned in our previous study (13) but not detected in synovial fluid derived RASF (Supplementary Table 2).

Of the 24 genes which were differentially expressed in both the synovial fluid and tissue derived RASF KD samples (Supplementary table 1) *NPTX1* and *MX1* were observed to be significantly differentially regulated. *NPTX1* was upregulated in both RASF KD samples and negligibly in OASF KD sample. *NPTX1* has been shown to be downregulated in RASF as compared to healthy SF in a microarray-based study (30) , which supports our present findings. On the other hand, *MX1* was upregulated in both RASF KD samples but down regulated in OASF. *MX1* is an interferon (IFN) response gene, and it has been shown to be correlated with disease activity in RASF. It has also been demonstrated that *MX1* may be an important gene for predicting the treatment response in RA patients (31).

Of the remaining 22 genes which were differentially regulated between RASF *ARL15* WT and KD, most significant was the downregulated *COMP*. This gene has been shown to play an important role for the stabilization of extracellular matrix. Higher amount of COMP has also shown to be correlated with severe disease in RA patients, implying its potential role as a biomarker for RA (32). Downregulation of *COMP* in both *ARL15* KD RASF samples in our study reinforce the role of *ARL15* in RA biology. In addition, *CTGF, PTX3, IFI6, USP18* and *CD248* are five other differentially regulated genes which seem to be functionally relevant for RA biology. *CTGF* has been shown to be involved in bone degradation of RA patients by activating osteoclastogenesis (33), and its inhibition in CIA mice model has been reported to ameliorate the disease (34) Our finding of *CTGF* being downregulated upon *ARL15* KD may yet again support the likely disease causal role of *ARL15*. *PTX3* has been reported to be overexpressed in RA cases (35). It has also been shown to be produced upon the activation of proinflammatory cytokines and is long thought to be involved in cardiovascular disease (CVD) prognosis (36). Of note, RA is associated with increased CVD risk (37) and *PTX3* was downregulated upon *ARL15* KD in both RA samples. These findings taken together amply support a likely role of *ARL15* in metabolic syndromes as well. On the other hand, *IFI6* and *USP18* are major IFN signaling genes. Inhibition of *USP18* has been correlated with inflammation (38) and upregulation of *IFI6* is observed in good responders of TNF antagonists (39) . Both *IFI6* and *USP18* were upregulated in *ARL15* KD RASF, yet again supporting a putative role of *ARL15* in RA. A previous study on *CD248* knockout mice showed less severe collagen induced arthritis compared to wild type with reduction in synovial hyperplasia and cartilage destruction (40) Our finding of reduced *CD248* expression upon *ARL15* KD further strengthens the druggability of *ARL15* in decreasing severity of RA.

MH7A cells: On analysis of transcriptome data in MH7A cells *ARL15* was the most significantly downregulated (Log 2 FC = -3.9, P-adjusted value = 1.39E-49) validated by qPCR (Figure 1C). Most of the differentially expressed genes in MH7A KD were informative (Supplementary table 5) but were notably different compared to RASF findings. In MH7A samples major cytokine and chemokine genes such as *IL1A, IL8, IL24, CXCL1, CXCL2,* and *CXCL3* were among the 12 significantly upregulated functionally relevant genes (Supplementary table 5) and validated through qPCR (Figure 1D) but are in stark contrast to expectation from a ARL15KD background. On the other hand, the nine downregulated genes included *MSN, TMED10, RARRES2, CD82, DOCK2, TGFB2, TLR4, FPR1* and *TFPI* (Figure 1C). Of these, *MSN* is part of the ERM (ezrin/radixin/moesin) proteins, where increased phosphorylation in response to cytokines promotes RASF proliferation (41). *TMED10*, acting as a receptor for ARF1-GDP, participates in COPI-vesicle formation on the Golgi membrane, enhancing coatomer-dependent GTPase-activating activity of ARFGAP2. *RARRES2* encodes chemerin, an adipokine involved in inflammation, adipogenesis, angiogenesis, and energy metabolism, also promoting adipocyte differentiation (42). *CD82* functions as a tumor metastasis suppressor; its overexpression in SFs reduces RASF migration and adhesion under pro-inflammatory stimuli (43). DOCK2 acts as a guanine exchange factor (GEF), mediating Rac1/2 activation and regulating immune cell migration (44) . TGFB2, an anti-inflammatory cytokine, is implicated in RA pathogenesis (45). TLR4 upregulates inflammatory cytokines and chemokines (IL-6, IL-8, and MMP3) in RASF (46). TFPI inhibits tissue factor (TF)-mediated inflammation in arthritis synovial joints (47).

At this juncture it is important to discuss the notable differences between patient derived cells and the cell line. A large number of genes were differentially expressed between RASF and MH7A control samples (Figure 3A). Notable differences in the transcriptome profiles were also observed between these two sample types (Figure3B) which was effectively captured in the correlation plot (Figure 3C). Conversely, as expected, samples derived from synovial tissue in the study showed a higher correlation with the Caucasian RA synovial tissue sample which was used for comparison. These findings reiterate the basic differences between patient derived primary cells and an established cell line. It may be important to mention here that FPKM values observed in the two RASF WT samples in our study, were comparable with the values reported for Caucasian RASF but not for healthy SF samples (48) (Supplementary Table 6 ) further supporting our findings.

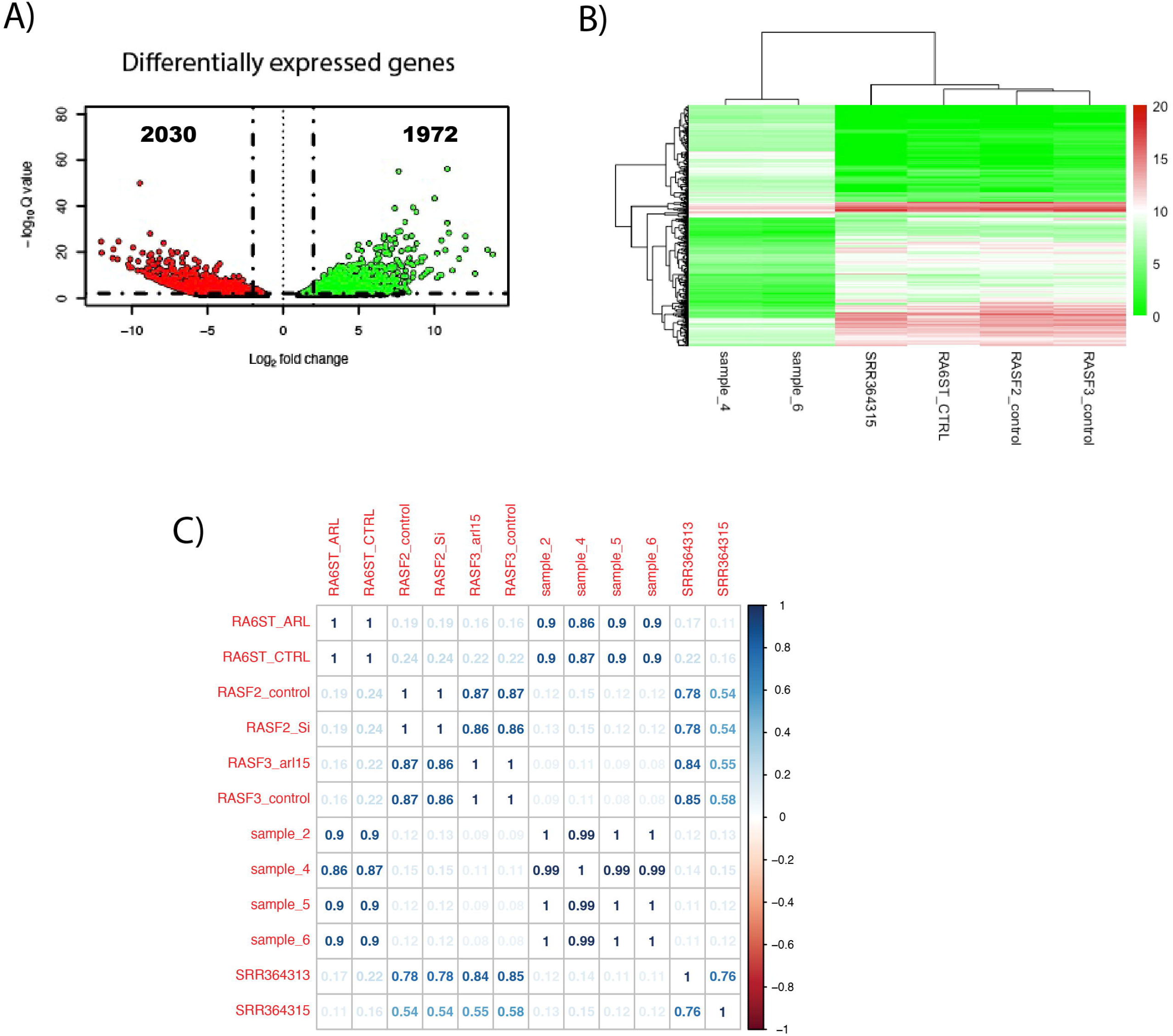

In addition, this study using two different experimental samples mainly patient derived RASF and MH7A cell line uncovers a basic fact on the differences in their utility to characterize a susceptibility gene(s) of interest. RASF in early passages seem to represent/capture the key players in active disease while the cell line largely manifests inflammation related markers. Such differences underscore the effect of immortalization on gene expression and consequent under representation of disease related markers. Nevertheless, both these study designs together seem to provide useful insights into the role of *ARL15* in RA.

In summary, this study on *ARL15* KD in RASF provides important insights into the role of this small GTPase in regulation of RA disease biology via maintenance of connective tissue architecture (*CTGF* and *COMP*); reduction of inflammation (*PTX3* and *USP18*); and increase in apoptosis; together with some leads from MH7A KD cells through cytokine pathways. Despite significant therapeutic advances, RA treatment remains a major challenge encouraging alternate approaches such as genomic medicine. In this context, taken together the leads from this study using a combination of *ex vivo* and *in vitro* models are suggestive of a likely regulatory role of *ARL15* in RA. They also highlight the need for further characterization of *ARL15* to explore its druggability using animal models of RA.

---

**Supplementary figure1:**
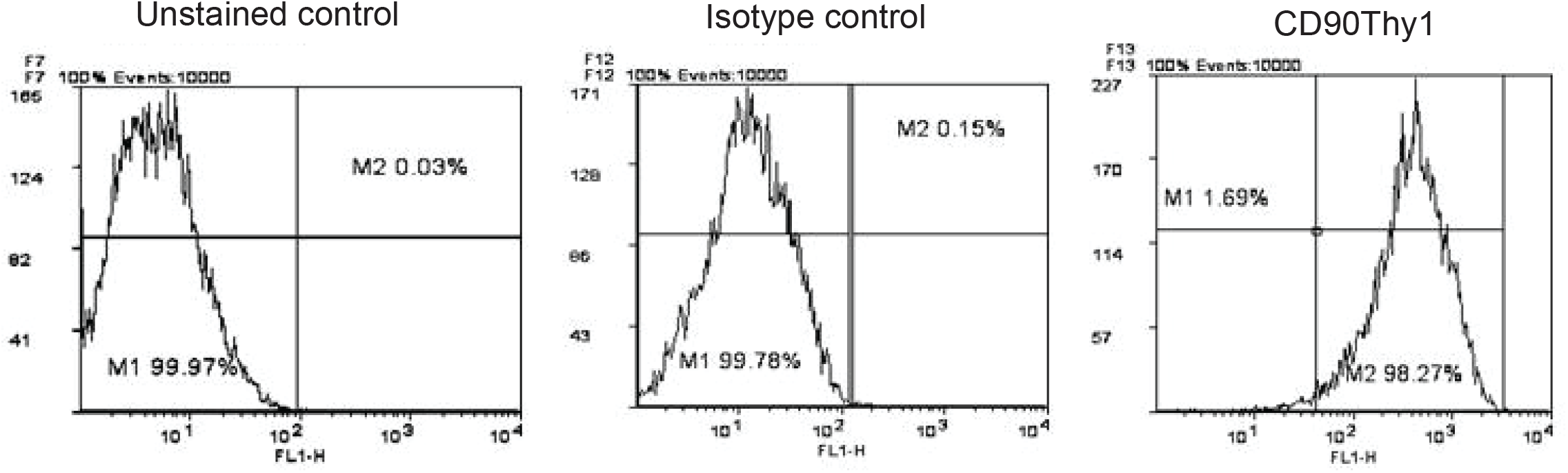
Confirmation of synovial fibroblast homogeniety by flow staining. From left to right synovial fibroblast cells were run on FACS caliber (BD) with unstained control followed by Isotype and CD90Thy1 (FITC) antibody.

**Supplementary figure2:**
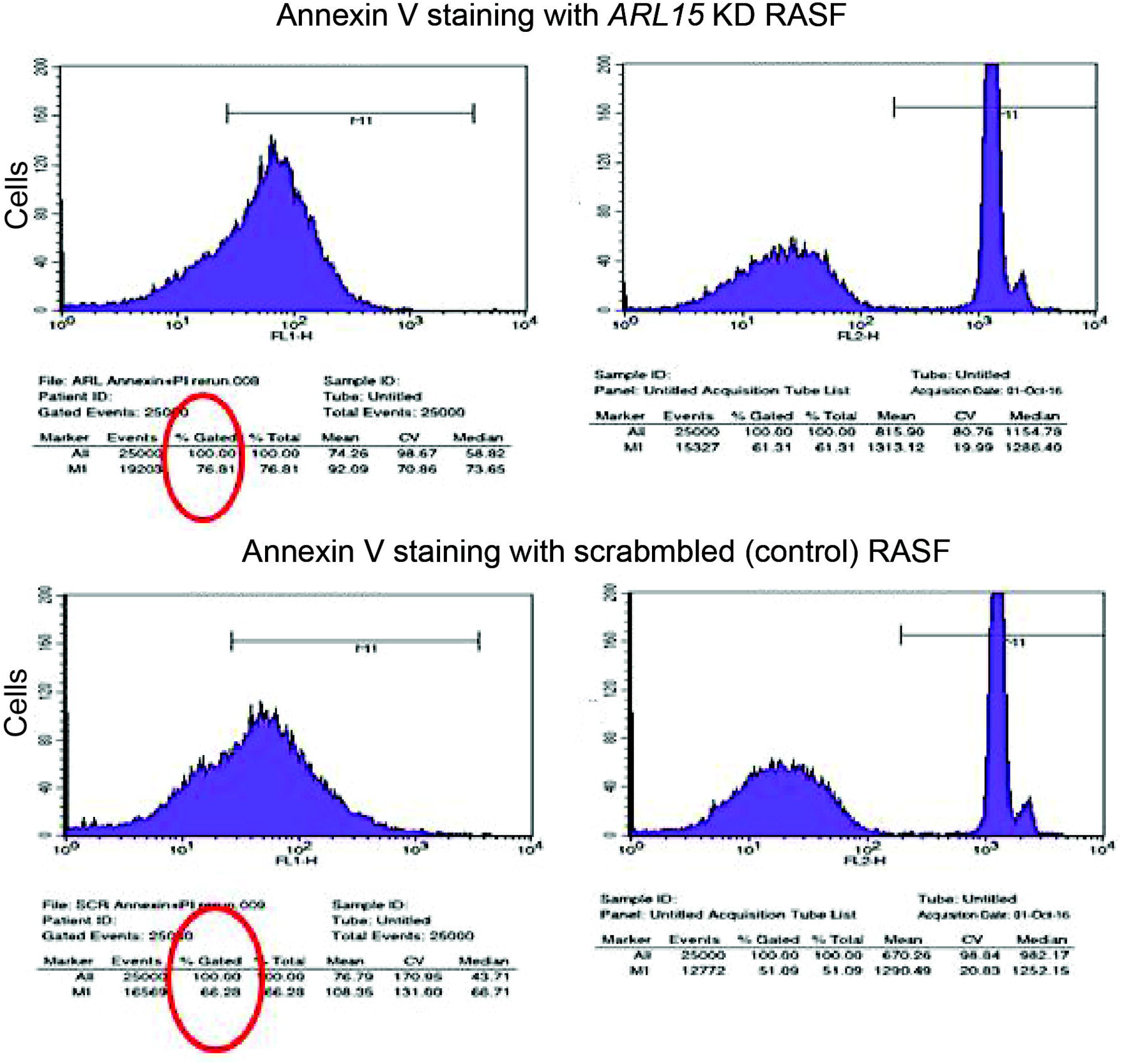
Annexin V staining and flow staining analysiswith Scrambled control RASF (Annexin + Pl run). Higher cell death upon ARL15 KD (76.81 %) as compared to the control (66.26%)

