## Supplementary table 1 for "Dissecting *ARL15* Function in Rheumatoid Arthritis: Insights from *Ex Vivo* and *In Vitro* Synovial Fibroblast Models"

**Supplementary table 1: Genes differentially expressed on *ARL15* KD and shared between RASF fluid derived and RASF tissue derived samples along with their expression in OASF control and KD groups. Top two genes namely, NPTX1, MX1 were common in all three samples, rest of the genes were common in RA samples only. Table is arranged based on the average P value for all the samples.**

| **Differentially regulated genes upon *ARL15* knock down** | | | | | | | | | | | | | | |
| --- | --- | --- | --- | --- | --- | --- | --- | --- | --- | --- | --- | --- | --- | --- |
| **No.** | **Gene name** | **RASF3 Ctrl FPKM** | **RASF3 ARL KD FPKM** | **log2(foldchange) _RA3** | **P_value_ (RA3)** | **RASF6 Ctrl FPKM** | **RASF6 ARL KD FPKM** | **log2(fold change) _RA6** | **P_value_ (RA6)** | **OASF1 Ctrl FPKM** | **OASF1 ARL KD FPKM** | **log2(fold change) _OA1** | **P_value_ (OA)** | **P_avg_** |
| **1** | **Neuronal pentraxin 1 (NPTX1)** | **10.65** | **23.22** | **1.12** | **5.00E-05** | **2.06** | **4.88** | **1.24** | **0.0013** | **0.70** | **1.45** | **1.04** | **0.0005** | **6.17E-04** |
| **2** | **Myxovirus resistance1 (MX1)** | **4.56** | **9.09** | **1.00** | **0.0058** | **0.78** | **4.22** | **2.43** | **0.0011** | **10.21** | **5.32** | **-0.94** | **0.0007** | **2.52E-03** |
| **3** | **Cartilage oligomatrix protein (COMP)** | **253.07** | **141.45** | **-0.84** | **0.00035** | **8.88** | **3.41** | **-1.38** | **0.0005** | **366.16** | **428.17** | **na** | **ns** | **4.25E-04** |
| **4** | **Connective tissue growth factor (CTGF)** | **434.45** | **271.29** | **-0.68** | **0.00645** | **117.39** | **64.86** | **-0.86** | **0.0044** | **454.01** | **473.67** | **na** | **ns** | **5.40E-03** |
| **5** | **Potassium voltage-gated channel subfamily D member 3(KCND3)** | **7.86** | **5.06** | **-0.64** | **0.014** | **0.88** | **0.26** | **-1.78** | **0.0023** | **0.46** | **0.40** | **na** | **ns** | **8.13E-03** |
| **6** | **Endothelial cell-specific molecule 1 (ESM1)** | **2.35** | **0.81** | **-1.54** | **0.01715** | **4.82** | **23.98** | **2.31** | **5E-05** | **77.32** | **68.53** | **na** | **ns** | **8.60E-03** |
| **7** | **Pentraxin (PTX3)** | **43.87** | **27.71** | **-0.66** | **0.0089** | **716.36** | **386.55** | **-0.89** | **0.0094** | **380.99** | **374.00** | **na** | **ns** | **9.15E-03** |
| **8** | **MCF.2 cell line derived transforming sequence-like (MCF2L)** | **2.63** | **5.58** | **1.09** | **0.0186** | **1.89** | **0.47** | **-1.99** | **0.0052** | **8.19** | **8.06** | **na** | **ns** | **1.19E-02** |
| **9** | **Leucine Rich Repeat Containing 17 (LRRC17)** | **15.27** | **22.41** | **0.55** | **0.02895** | **3.95** | **11.99** | **1.60** | **0.0002** | **1.57** | **1.42** | **na** | **ns** | **1.46E-02** |
| **10** | **(Follistatin-like 3) FSTL3** | **20.76** | **12.02** | **-0.79** | **0.0054** | **15.73** | **9.13** | **-0.79** | **0.0279** | **32.84** | **33.62** | **na** | **ns** | **1.67E-02** |
| **11** | **Keratin 19 (KRT19)** | **28.93** | **16.64** | **-0.80** | **0.003** | **3.40** | **6.39** | **0.91** | **0.0307** | **19.80** | **19.73** | **na** | **ns** | **1.69E-02** |
| **12** | **Proline And Arginine Rich End Leucine Rich Repeat Protein (PRELP)** | **13.92** | **8.63** | **-0.69** | **0.0045** | **1.64** | **0.86** | **-0.93** | **0.0294** | **90.10** | **88.89** | **na** | **ns** | **1.70E-02** |
| **13** | **Aggrecan (ACAN)** | **23.19** | **9.96** | **-1.22** | **0.0071** | **50.61** | **29.81** | **-0.76** | **0.0284** | **71.87** | **59.79** | **na** | **ns** | **1.78E-02** |
| **14** | **Endosialin (CD248)** | **215.26** | **153.80** | **-0.48** | **0.0387** | **32.53** | **14.04** | **-1.21** | **5E-05** | **531.79** | **471.48** | **na** | **ns** | **1.94E-02** |
| **15** | **Cbp/p300 interacting transactivator with Glu/Asp rich carboxy-terminal domain 2 (CITED2)** | **197.19** | **141.95** | **-0.47** | **0.0381** | **68.60** | **34.04** | **-1.01** | **0.001** | **70.29** | **67.16** | **na** | **ns** | **1.95E-02** |
| **16** | **Membrane-associated transporter protein (SLC45A2)** | **4.71** | **7.01** | **0.58** | **0.0392** | **0.46** | **1.38** | **1.58** | **0.002** | **8.13** | **8.26** | **na** | **ns** | **2.06E-02** |
| **17** | **Ubiquitin-specific peptidase 18 (USP18)** | **0.82** | **1.92** | **1.23** | **0.02685** | **1.52** | **3.29** | **1.11** | **0.0176** | **4.16** | **3.31** | **na** | **ns** | **2.22E-02** |
| **18** | **Collagen type XIV alpha 1 chain (COL14A1)** | **45.40** | **77.59** | **0.77** | **0.00095** | **13.17** | **19.87** | **0.59** | **0.0442** | **81.61** | **86.99** | **na** | **ns** | **2.26E-02** |
| **19** | **Centromere Protein K (CENPK)** | **0.21** | **1.71** | **2.99** | **0.00245** | **18.78** | **30.14** | **0.68** | **0.0482** | **5.28** | **4.36** | **na** | **ns** | **2.53E-02** |
| **20** | **Nerve Growth Factor (NGF)** | **12.38** | **7.70** | **-0.69** | **0.0377** | **13.87** | **7.63** | **-0.86** | **0.0237** | **1.26** | **1.03** | **na** | **ns** | **3.07E-02** |
| **21** | **G Protein-Coupled Receptor 153 (GPR153)** | **8.91** | **13.11** | **0.56** | **0.0245** | **3.39** | **5.35** | **0.66** | **0.0479** | **2.15** | **2.34** | **na** | **ns** | **3.62E-02** |
| **22** | **Interleukin 7 Receptor (IL7R)** | **1.22** | **0.38** | **-1.69** | **0.0368** | **2.63** | **5.10** | **0.96** | **0.0391** | **2.28** | **2.32** | **na** | **ns** | **3.80E-02** |
| **23** | **Interferon alpha inducible protein 6 (IFI6)** | **68.12** | **119.7** | **0.81309** | **8.00E-04** | **25.75** | **46.05** | **0.83852** | **0.0087** | **31.91** | **30.76** | **na** | **ns** | **4.73E-03** |
| **24** | **Sarcolipin (SLN)** | **1.18707** | **0** | **NA** | **0.01095** | **0** | **0.57629** | **INF** | **0.01835** | **0** | **0** | **na** | **ns** | **0.01465** |
| **25** | **Glutamate Receptor Interacting Protein 1 (GRIP1)** | **0.18945** | **2.38448** | **3.65378** | **0.02455** | **0.54842** | **0.07673** | **-2.83742** | **0.0283** | **0** | **0.05987** | **na** | **ns** | **0.026425** |
