## Supplementary table2 for "Dissecting *ARL15* Function in Rheumatoid Arthritis: Insights from *Ex Vivo* and *In Vitro* Synovial Fibroblast Models"

| **Supplementary table 2: 217 differentially regulated genes (P<0.05) obtained upon *ARL15* knockdown in synovial fluid derived RASF3** | | | | | |
| --- | --- | --- | --- | --- | --- |
| **Gene** | **Locus** | **RASF3 CONTROL FPKM** | **RASF3 (ARL15_siRNA)FPKM** | **log2(fold_change)** | **p_value** |
| **FER1L6** | **chr8:124864226-125183763** | **1.62** | **3.83** | **1.24** | **0.00005** |
| **HMOX1** | **chr22:35776353-35790207** | **32.74** | **69.16** | **1.08** | **0.00005** |
| **KRT16** | **chr17:39766029-39772151** | **35.95** | **14.76** | **-1.28** | **0.00005** |
| **MAFB** | **chr20:39314487-39317880** | **1.37** | **4.30** | **1.65** | **0.00005** |
| **NPTX1** | **chr17:78440947-78451643** | **10.65** | **23.22** | **1.12** | **0.00005** |
| **RNF144A** | **chr2:7052407-7218011** | **4.69** | **10.54** | **1.17** | **0.0001** |
| **C11orf87** | **chr11:109225811-109454633** | **14.42** | **7.87** | **-0.87** | **0.00015** |
| **MARCKSL1** | **chr1:32799432-32801980** | **14.03** | **27.74** | **0.98** | **0.0002** |
| **AGT** | **chr1:230838268-230937749** | **3.46** | **8.02** | **1.21** | **0.00035** |
| **COMP** | **chr19:18893582-18902123** | **253.07** | **141.45** | **-0.84** | **0.00035** |
| **TLL2** | **chr10:98124362-98273675** | **2.38** | **0.88** | **-1.44** | **0.00045** |
| **ABCA1** | **chr9:107543282-107691173** | **10.55** | **18.10** | **0.78** | **0.0005** |
| **IFI6** | **chr1:27992571-27998729** | **68.12** | **119.68** | **0.81** | **0.0008** |
| **SOX4** | **chr6:21593971-21598847** | **17.33** | **29.44** | **0.76** | **0.0009** |
| **COL14A1** | **chr8:121072018-121384275** | **45.40** | **77.59** | **0.77** | **0.00095** |
| **GEM** | **chr8:95261480-95274578** | **24.68** | **42.77** | **0.79** | **0.0017** |
| **KIF26B** | **chr1:245318286-245872733** | **3.28** | **5.73** | **0.81** | **0.00175** |
| **EFEMP1** | **chr2:56093101-56151274** | **258.00** | **154.05** | **-0.74** | **0.0018** |
| **KRT14** | **chr17:39738530-39743173** | **45.00** | **25.29** | **-0.83** | **0.00185** |
| **HHAT** | **chr1:210501595-210849638** | **2.16** | **4.47** | **1.05** | **0.00215** |
| **CENPK** | **chr5:64813592-64858998** | **0.21** | **1.71** | **2.99** | **0.00245** |
| **NOV** | **chr8:120428545-120474590** | **78.35** | **48.70** | **-0.69** | **0.00275** |
| **KRT19** | **chr17:39679868-39684560** | **28.93** | **16.64** | **-0.80** | **0.003** |
| **PDE4D** | **chr5:58264864-59843484** | **0.46** | **3.07** | **2.75** | **0.0032** |
| **LGR5** | **chr12:71518864-71980090** | **19.27** | **10.62** | **-0.86** | **0.00325** |
| **TNXB** | **chr6:32006041-32096030** | **16.14** | **8.67** | **-0.90** | **0.00335** |
| **FAM134B** | **chr5:16473146-16630078** | **2.31** | **5.24** | **1.18** | **0.0036** |
| **KCTD12** | **chr13:77454311-77460540** | **9.05** | **14.44** | **0.67** | **0.004** |
| **SLIT3** | **chr5:168081517-168728133** | **29.09** | **17.76** | **-0.71** | **0.00405** |
| **MEX3B** | **chr15:82334118-82338482** | **3.32** | **5.91** | **0.83** | **0.00435** |
| **SCD** | **chr10:102106880-102124591** | **19.89** | **31.88** | **0.68** | **0.00445** |
| **PRELP** | **chr1:203444955-203460480** | **13.92** | **8.63** | **-0.69** | **0.0045** |
| **PINX1** | **chr8:10581277-10697394** | **2.78** | **6.80** | **1.29** | **0.0048** |
| **SCRG1** | **chr4:174305851-174327531** | **2.55** | **1.15** | **-1.15** | **0.00485** |
| **PPP1R13B** | **chr14:104200088-104313927** | **3.42** | **0.97** | **-1.82** | **0.0049** |
| **FGGY** | **chr1:59762309-60254854** | **7.46** | **3.11** | **-1.26** | **0.00495** |
| **CCDC169** | **chr13:36742344-36871979** | **0.14** | **1.37** | **3.31** | **0.0052** |
| **TMEM117** | **chr12:44229769-44783545** | **5.50** | **10.98** | **1.00** | **0.00535** |
| **FSTL3** | **chr19:676391-683385** | **20.76** | **12.02** | **-0.79** | **0.0054** |
| **CLEC3B** | **chr3:44956748-45077563** | **1564.05** | **916.56** | **-0.77** | **0.0058** |
| **MX1** | **chr21:42792230-42831141** | **4.56** | **9.09** | **1.00** | **0.0058** |
| **BAALC** | **chr8:104133259-104345094** | **19.57** | **42.50** | **1.12** | **0.00615** |
| **NOVA1** | **chr14:26912298-27066960** | **1.05** | **0.03** | **-5.26** | **0.00625** |
| **CTGF** | **chr6:132269315-132398533** | **434.45** | **271.29** | **-0.68** | **0.00645** |
| **GGT5** | **chr22:24615621-24641110** | **0.42** | **2.13** | **2.35** | **0.0067** |
| **TTF2** | **chr1:117602924-117650075** | **4.37** | **1.17** | **-1.90** | **0.0069** |
| **GMPR** | **chr6:16238810-16295780** | **9.56** | **16.36** | **0.78** | **0.007** |
| **RHOB** | **chr2:20646834-20649200** | **59.96** | **38.54** | **-0.64** | **0.00705** |
| **ACAN** | **chr15:89346673-89418585** | **23.19** | **9.96** | **-1.22** | **0.0071** |
| **MDK** | **chr11:46402305-46405375** | **40.57** | **65.61** | **0.69** | **0.0071** |
| **ADAMTS14** | **chr10:72432558-72522197** | **6.52** | **10.41** | **0.67** | **0.00735** |
| **ZNF414** | **chr19:8575461-8579048** | **11.68** | **5.44** | **-1.10** | **0.00755** |
| **CCL2** | **chr17:32581899-32584222** | **10.17** | **4.38** | **-1.22** | **0.008** |
| **ITPKB** | **chr1:226819390-226927024** | **3.51** | **5.76** | **0.71** | **0.0081** |
| **ZNF287** | **chr17:16454700-16472520** | **7.21** | **2.73** | **-1.40** | **0.0082** |
| **PLXDC1** | **chr17:37213271-37310647** | **14.04** | **26.59** | **0.92** | **0.00845** |
| **GPNMB** | **chr7:23275585-23314727** | **268.35** | **415.00** | **0.63** | **0.0088** |
| **PTX3** | **chr3:156893011-157251408** | **43.87** | **27.71** | **-0.66** | **0.0089** |
| **RGS17** | **chr6:153325593-153452384** | **0.94** | **2.19** | **1.22** | **0.0091** |
| **STRC** | **chr15:43885251-44010458** | **1.03** | **0.06** | **-4.13** | **0.0096** |
| **ELN** | **chr7:73442118-73484237** | **94.79** | **61.91** | **-0.61** | **0.00965** |
| **CRIP1** | **chr14:105952653-105965912** | **233.19** | **149.84** | **-0.64** | **0.0102** |
| **CLCA2** | **chr1:86889768-86922241** | **1.38** | **2.99** | **1.12** | **0.01025** |
| **PEG10** | **chr7:94285636-94299007** | **3.02** | **4.97** | **0.72** | **0.01025** |
| **TMEM35** | **chrX:100264334-100351353** | **7.15** | **13.00** | **0.86** | **0.01055** |
| **CRYM** | **chr16:21244985-21329912** | **2.02** | **6.67** | **1.72** | **0.0106** |
| **PTCHD4** | **chr6:47845763-48036425** | **0.68** | **1.63** | **1.26** | **0.0107** |
| **EEPD1** | **chr7:36192757-36341152** | **4.22** | **9.89** | **1.23** | **0.01075** |
| **SLN** | **chr11:107578103-107595135** | **1.19** | **0.00** | **#NAME?** | **0.01095** |
| **SOCS2** | **chr12:93936238-93977263** | **9.67** | **5.21** | **-0.89** | **0.011** |
| **TRIM45** | **chr1:117653681-117665209** | **1.38** | **3.22** | **1.22** | **0.0111** |
| **PTGES** | **chr9:132500609-132515326** | **14.81** | **9.08** | **-0.71** | **0.01125** |
| **MYBL1** | **chr8:67474409-67526482** | **4.47** | **2.16** | **-1.05** | **0.0114** |
| **SCUBE2** | **chr11:9025708-9159661** | **1.32** | **3.66** | **1.47** | **0.01165** |
| **ARHGAP24** | **chr4:86396266-86923823** | **9.65** | **15.88** | **0.72** | **0.0117** |
| **CDH4** | **chr20:59827481-60515673** | **1.07** | **0.52** | **-1.06** | **0.01185** |
| **CLIC2** | **chrX:154505499-154563966** | **1.13** | **3.31** | **1.55** | **0.01185** |
| **CLIC3** | **chr9:139889086-139891255** | **14.29** | **7.74** | **-0.88** | **0.01245** |
| **APITD1** | **chr1:10489937-10512210** | **19.25** | **10.77** | **-0.84** | **0.0132** |
| **KCND3** | **chr1:112313283-112531777** | **7.86** | **5.06** | **-0.64** | **0.014** |
| **GRB14** | **chr2:165349321-165478358** | **0.53** | **2.44** | **2.21** | **0.01405** |
| **SEC16B** | **chr1:177893090-178007142** | **1.28** | **5.27** | **2.05** | **0.01405** |
| **CES4A** | **chr16:67022491-67043661** | **5.67** | **2.69** | **-1.08** | **0.0145** |
| **CRABP2** | **chr1:156669397-156675608** | **186.62** | **125.98** | **-0.57** | **0.0154** |
| **ATHL1** | **chr11:289134-296107** | **11.36** | **6.75** | **-0.75** | **0.0164** |
| **LRRC32** | **chr11:76368099-76381791** | **5.87** | **10.77** | **0.88** | **0.01645** |
| **ZNF674** | **chrX:46357161-46404892** | **2.37** | **0.77** | **-1.62** | **0.01675** |
| **PRKCH** | **chr14:61654276-62125414** | **2.66** | **0.79** | **-1.76** | **0.0168** |
| **SYNPO** | **chr5:149980641-150038782** | **2.81** | **1.52** | **-0.89** | **0.01695** |
| **UAP1L1** | **chr9:139971952-139978991** | **6.97** | **10.70** | **0.62** | **0.0171** |
| **ESM1** | **chr5:54273691-54338672** | **2.35** | **0.81** | **-1.54** | **0.01715** |
| **BMP2** | **chr20:6748310-6760927** | **1.28** | **2.40** | **0.90** | **0.01735** |
| **FXYD3** | **chr19:35606731-35615228** | **0.30** | **1.37** | **2.19** | **0.0175** |
| **PPAP2C** | **chr19:281039-291393** | **1.36** | **3.68** | **1.44** | **0.01775** |
| **NEDD9** | **chr6:11173684-11382581** | **13.35** | **9.07** | **-0.56** | **0.01795** |
| **ANKUB1** | **chr3:149478891-149942977** | **1.21** | **0.00** | **#NAME?** | **0.0184** |
| **MCF2L** | **chr13:113548691-113754053** | **2.63** | **5.58** | **1.09** | **0.0186** |
| **TNFRSF11B** | **chr8:119935795-119964439** | **53.28** | **34.07** | **-0.64** | **0.0187** |
| **MN1** | **chr22:28144264-28197486** | **10.21** | **6.93** | **-0.56** | **0.01885** |
| **STMN1** | **chr1:26210671-26233482** | **91.44** | **135.82** | **0.57** | **0.0189** |
| **COL15A1** | **chr9:101705460-101833069** | **4.36** | **2.62** | **-0.74** | **0.019** |
| **PAMR1** | **chr11:35453369-35551848** | **107.24** | **73.94** | **-0.54** | **0.0193** |
| **MYEOV** | **chr11:69061604-69182494** | **0.78** | **3.17** | **2.03** | **0.02015** |
| **CACNB2** | **chr10:18429605-18948214** | **1.31** | **0.26** | **-2.33** | **0.021** |
| **POT1** | **chr7:124462439-125019375** | **4.90** | **9.47** | **0.95** | **0.021** |
| **SEMA3B** | **chr3:50304989-50314977** | **33.91** | **21.85** | **-0.63** | **0.021** |
| **PLEKHO1** | **chr1:150121372-150136916** | **37.32** | **58.34** | **0.64** | **0.02115** |
| **CORO2B** | **chr15:68871307-69020145** | **4.23** | **6.59** | **0.64** | **0.02125** |
| **ARAP3** | **chr5:141032967-141061788** | **0.99** | **2.11** | **1.09** | **0.02135** |
| **MYO15B** | **chr17:73584138-73704142** | **3.05** | **0.83** | **-1.88** | **0.0215** |
| **RGMA** | **chr15:93586635-93632433** | **2.45** | **4.55** | **0.89** | **0.022** |
| **IGDCC4** | **chr15:65673801-65715410** | **0.90** | **2.15** | **1.26** | **0.02205** |
| **POLR3F** | **chr20:18364010-18465287** | **4.07** | **8.43** | **1.05** | **0.0231** |
| **LAMB3** | **chr1:209788214-209825811** | **1.32** | **2.86** | **1.11** | **0.0232** |
| **KRT8** | **chr12:53290976-53346686** | **0.35** | **1.60** | **2.18** | **0.02335** |
| **IL6** | **chr7:22765013-22771621** | **5.83** | **2.41** | **-1.27** | **0.0243** |
| **GPR153** | **chr1:6307405-6321035** | **8.91** | **13.11** | **0.56** | **0.0245** |
| **GRIP1** | **chr12:66741210-67197966** | **0.19** | **2.38** | **3.65** | **0.02455** |
| **LARP1B** | **chr4:128982422-129144086** | **6.60** | **4.13** | **-0.68** | **0.0247** |
| **CTSK** | **chr1:150768683-150780799** | **567.02** | **843.09** | **0.57** | **0.0253** |
| **TEAD2** | **chr19:49834873-49891338** | **9.46** | **16.24** | **0.78** | **0.02645** |
| **PDGFD** | **chr11:103777913-104035107** | **1.30** | **2.43** | **0.91** | **0.02665** |
| **THOC3** | **chr5:175333941-175461683** | **8.49** | **4.81** | **-0.82** | **0.02675** |
| **USP18** | **chr22:18632665-18660164** | **0.82** | **1.92** | **1.23** | **0.02685** |
| **DTWD2** | **chr5:118173016-118324240** | **1.79** | **0.94** | **-0.93** | **0.027** |
| **RIN3** | **chr14:92980117-93155339** | **9.29** | **5.30** | **-0.81** | **0.02705** |
| **TUFT1** | **chr1:151512780-151556059** | **14.49** | **9.30** | **-0.64** | **0.02705** |
| **PDXP** | **chr22:38030660-38062941** | **5.15** | **2.41** | **-1.09** | **0.0271** |
| **CCDC171** | **chr9:15552894-16061661** | **2.08** | **0.64** | **-1.69** | **0.02725** |
| **GALM** | **chr2:38893051-38968379** | **9.45** | **14.41** | **0.61** | **0.0277** |
| **FZD1** | **chr7:90893782-90898123** | **42.73** | **30.07** | **-0.51** | **0.02775** |
| **TNNC2** | **chr20:44451852-44462384** | **1.84** | **4.81** | **1.39** | **0.0279** |
| **KRT34** | **chr17:39533901-39538655** | **2.36** | **1.01** | **-1.22** | **0.0281** |
| **PSAT1** | **chr9:80912058-80945009** | **19.17** | **13.06** | **-0.55** | **0.02885** |
| **LRRC17** | **chr7:102453307-102715286** | **15.27** | **22.41** | **0.55** | **0.02895** |
| **NCAM1** | **chr11:112830001-113149158** | **1.05** | **0.31** | **-1.75** | **0.02905** |
| **CLGN** | **chr4:141309608-141349122** | **0.62** | **1.50** | **1.27** | **0.02945** |
| **KLHDC9** | **chr1:161068150-161070136** | **4.30** | **1.93** | **-1.16** | **0.03025** |
| **MARK2** | **chr11:63606399-63678491** | **5.01** | **8.18** | **0.71** | **0.03075** |
| **ARL15** | **chr5:53179774-53606412** | **6.21** | **3.13** | **-0.99** | **0.0309** |
| **KRT7** | **chr12:52626303-52702947** | **5.93** | **2.37** | **-1.33** | **0.0314** |
| **SPINT2** | **chr19:38734674-38795649** | **3.01** | **0.63** | **-2.26** | **0.0314** |
| **EDNRA** | **chr4:148402068-148466106** | **1.64** | **3.23** | **0.98** | **0.03185** |
| **TMEM132B** | **chr12:125671381-126146917** | **1.85** | **5.54** | **1.59** | **0.0319** |
| **RBPMS2** | **chr15:65032090-65067786** | **1.66** | **0.67** | **-1.32** | **0.03195** |
| **FOLR3** | **chr11:71825914-71850936** | **1.63** | **6.89** | **2.08** | **0.032** |
| **DBNDD2** | **chr20:43990576-44039250** | **33.33** | **19.97** | **-0.74** | **0.03285** |
| **IGF1** | **chr12:102789644-102874423** | **1.18** | **0.08** | **-3.92** | **0.03335** |
| **S1PR1** | **chr1:101702443-101707074** | **2.75** | **4.72** | **0.78** | **0.03355** |
| **HCLS1** | **chr3:121350245-121379774** | **0.38** | **1.40** | **1.86** | **0.0338** |
| **SFRP4** | **chr7:37723398-38065297** | **10.51** | **4.64** | **-1.18** | **0.03395** |
| **FAM231D** | **chr1:149646784-149719362** | **1.11** | **0.21** | **-2.41** | **0.03435** |
| **EVI5** | **chr1:92974252-93257961** | **10.18** | **5.56** | **-0.87** | **0.0346** |
| **SEPP1** | **chr5:42756902-42887494** | **1.84** | **3.70** | **1.01** | **0.03465** |
| **POLR3D** | **chr8:22102616-22132675** | **5.43** | **8.94** | **0.72** | **0.03475** |
| **PTS** | **chr11:112097087-112140678** | **20.68** | **13.01** | **-0.67** | **0.035** |
| **ERCC6** | **chr10:50627322-50747584** | **9.06** | **5.44** | **-0.73** | **0.03525** |
| **ZNF718** | **chr4:123965-157779** | **1.91** | **3.70** | **0.96** | **0.0353** |
| **AURKB** | **chr17:8108055-8113918** | **1.68** | **0.45** | **-1.89** | **0.03535** |
| **APOBEC3C** | **chr22:39410087-39429281** | **26.03** | **37.31** | **0.52** | **0.0354** |
| **DYNLRB2** | **chr16:80099732-80597032** | **0.13** | **1.68** | **3.73** | **0.03565** |
| **HAPLN3** | **chr15:89420518-89438857** | **46.53** | **31.74** | **-0.55** | **0.0357** |
| **PLIN2** | **chr9:19108372-19149288** | **47.75** | **66.65** | **0.48** | **0.03595** |
| **SPP1** | **chr4:88896818-88904562** | **0.62** | **1.94** | **1.64** | **0.03595** |
| **LEPREL1** | **chr3:189674516-189862635** | **3.72** | **1.44** | **-1.37** | **0.03615** |
| **C20orf96** | **chr20:251503-271390** | **3.85** | **6.44** | **0.74** | **0.03665** |
| **ARHGEF37** | **chr5:148931509-149014531** | **0.44** | **1.51** | **1.78** | **0.03675** |
| **IL7R** | **chr5:35852796-35879705** | **1.22** | **0.38** | **-1.69** | **0.0368** |
| **ZFP36** | **chr19:39897452-39900052** | **41.34** | **28.97** | **-0.51** | **0.0368** |
| **TMEM133** | **chr11:100862810-100864663** | **0.72** | **1.79** | **1.32** | **0.0369** |
| **FAM13C** | **chr10:60936349-61122939** | **2.46** | **4.58** | **0.89** | **0.0369** |
| **PRKAG2** | **chr7:151253196-151576299** | **8.89** | **5.35** | **-0.73** | **0.037** |
| **TBX5** | **chr12:114791735-114850636** | **1.30** | **3.21** | **1.30** | **0.037** |
| **FAM21A** | **chr10:51827647-51893269** | **5.91** | **10.65** | **0.85** | **0.0375** |
| **MGLL** | **chr3:127407908-127542051** | **45.77** | **30.88** | **-0.57** | **0.03755** |
| **KIF27** | **chr9:86444570-86536342** | **1.30** | **0.40** | **-1.69** | **0.0376** |
| **MAP2** | **chr2:210288781-210598842** | **0.33** | **1.21** | **1.87** | **0.0376** |
| **NGF** | **chr1:115825654-115910693** | **12.38** | **7.70** | **-0.69** | **0.0377** |
| **EPAS1** | **chr2:46520805-46613836** | **27.25** | **37.85** | **0.47** | **0.03775** |
| **CITED2** | **chr6:139693392-139695757** | **197.19** | **141.95** | **-0.47** | **0.0381** |
| **CD248** | **chr11:66080323-66086708** | **215.26** | **153.80** | **-0.48** | **0.0387** |
| **FBLN2** | **chr3:13573823-13679922** | **71.38** | **51.51** | **-0.47** | **0.03895** |
| **AIM1** | **chr6:106959729-107018326** | **4.71** | **7.01** | **0.58** | **0.0392** |
| **P2RX7** | **chr12:121570621-121623876** | **0.35** | **1.08** | **1.64** | **0.03935** |
| **TRIM47** | **chr17:73870241-73875627** | **28.72** | **17.22** | **-0.74** | **0.03945** |
| **NAF1** | **chr4:164029936-164088073** | **6.35** | **3.73** | **-0.77** | **0.03995** |
| **ZNF273** | **chr7:64330549-64391344** | **1.28** | **2.62** | **1.03** | **0.04035** |
| **ATP6V0E2** | **chr7:149564785-149577784** | **16.00** | **10.56** | **-0.60** | **0.0405** |
| **RBM19** | **chr12:114254542-114404176** | **4.78** | **7.84** | **0.71** | **0.0405** |
| **EGR2** | **chr10:64571755-64679660** | **5.48** | **8.08** | **0.56** | **0.04085** |
| **FOXRED2** | **chr22:36883236-36903148** | **3.97** | **6.02** | **0.60** | **0.04145** |
| **VWCE** | **chr11:61025761-61062896** | **11.32** | **17.86** | **0.66** | **0.04155** |
| **PORCN** | **chrX:48367349-48379202** | **20.47** | **13.73** | **-0.58** | **0.0418** |
| **ZNF131** | **chr5:43014515-43192123** | **11.29** | **18.94** | **0.75** | **0.0418** |
| **PCDH1** | **chr5:141232937-141258811** | **0.69** | **1.59** | **1.20** | **0.04205** |
| **MSRA** | **chr8:9911777-10286401** | **6.67** | **10.78** | **0.69** | **0.0425** |
| **TMEM60** | **chr7:77423044-77427897** | **13.33** | **20.19** | **0.60** | **0.04285** |
| **C14orf28** | **chr14:45366497-45381066** | **5.35** | **3.24** | **-0.72** | **0.0435** |
| **OVGP1** | **chr1:111956935-111970399** | **1.75** | **3.64** | **1.06** | **0.04385** |
| **ZNF341** | **chr20:32316736-32398903** | **0.56** | **1.15** | **1.03** | **0.04445** |
| **LEKR1** | **chr3:156543269-156763918** | **0.28** | **1.29** | **2.18** | **0.0445** |
| **TLR5** | **chr1:223282747-223316624** | **0.15** | **1.02** | **2.80** | **0.0457** |
| **ADAM32** | **chr8:38964508-39142430** | **0.86** | **3.03** | **1.82** | **0.04645** |
| **ALS2CL** | **chr3:46710486-46735194** | **4.62** | **2.76** | **-0.75** | **0.0465** |
| **CA12** | **chr15:63613576-63674360** | **47.44** | **66.94** | **0.50** | **0.0467** |
| **DNMT3A** | **chr2:25455844-25565459** | **2.82** | **5.31** | **0.92** | **0.04685** |
| **MAFK** | **chr7:1570349-1600457** | **24.50** | **17.10** | **-0.52** | **0.047** |
| **POLA2** | **chr11:65029232-65073060** | **6.66** | **3.60** | **-0.89** | **0.04705** |
| **CYTIP** | **chr2:158271130-158345473** | **0.35** | **1.51** | **2.11** | **0.04725** |
| **GTF2F1** | **chr19:6379579-6412415** | **36.46** | **24.35** | **-0.58** | **0.04755** |
| **CACNB4** | **chr2:152689289-152965716** | **4.17** | **2.19** | **-0.93** | **0.04815** |
| **NRN1** | **chr6:5998231-6007200** | **0.97** | **2.16** | **1.16** | **0.0485** |
| **JMJD7-PLA2G4B** | **chr15:42120282-42190773** | **1.36** | **2.72** | **1.00** | **0.0486** |
| **MKX** | **chr10:27961803-28056723** | **47.50** | **34.19** | **-0.47** | **0.04865** |
| **ZNF599** | **chr19:35248980-35264134** | **2.95** | **5.39** | **0.87** | **0.04875** |
| **ANKRD1** | **chr10:92671852-92681033** | **1.72** | **0.78** | **-1.14** | **0.04955** |
| **MKKS** | **chr20:10381656-10414870** | **10.14** | **14.41** | **0.51** | **0.0498** |
