## Supplementary table 3 for "Dissecting *ARL15* Function in Rheumatoid Arthritis: Insights from *Ex Vivo* and *In Vitro* Synovial Fibroblast Models"

| **Supplementary table 3: 340 differentially regulated genes (P<0.05) upon *ARL15* knockdown in synovial tissue derived RASF6** | | | | | |
| --- | --- | --- | --- | --- | --- |
| **Gene** | **locus** | **RASF6**  **CTRL FPKM** | **RASF6 (ARL15_siRNA)FPKM** | **log2(fold_change)** | **P_value** |
| **CD248** | **11:66080323-66086708** | **32.5272** | **14.0359** | **-1.21253** | **0.00005** |
| **ESM1** | **5:54273691-54338672** | **4.82356** | **23.9829** | **2.31383** | **0.00005** |
| **IL13RA2** | **X:114238537-114254540** | **51.809** | **125.761** | **1.27941** | **0.00005** |
| **ISG15** | **1:948802-949920** | **20.744** | **61.1526** | **1.55972** | **0.00005** |
| **KRT7,KRT86** | **12:52626303-52702947** | **84.7029** | **29.4077** | **-1.52621** | **0.00005** |
| **MMP1** | **11:102617698-102714534** | **5.62226** | **19.5662** | **1.79914** | **0.00005** |
| **NTS** | **12:86268072-86276770** | **0** | **1.25878** | **inf** | **0.00005** |
| **PHLDA1** | **12:76419226-76427712** | **4.13424** | **11.1182** | **1.42722** | **0.00005** |
| **PLAT** | **8:42032235-42065242** | **19.4527** | **66.0713** | **1.76405** | **0.00005** |
| **SYNPO2** | **4:119809995-119982402** | **40.892** | **16.9755** | **-1.26837** | **0.00005** |
| **THBD** | **20:23026269-23030378** | **6.08571** | **16.5452** | **1.44292** | **0.00005** |
| **THBS1** | **15:39872729-39891667** | **255.353** | **107.455** | **-1.24877** | **0.0001** |
| **IFI44** | **1:79115460-79129763** | **7.41683** | **22.0906** | **1.57456** | **0.00015** |
| **LRRC17** | **7:102453307-102715286** | **3.95461** | **11.994** | **1.6007** | **0.00015** |
| **MGAT4C** | **12:86372515-87232775** | **0** | **0.587542** | **inf** | **0.00015** |
| **DUSP6** | **12:89741008-89747048** | **7.76762** | **23.2756** | **1.58327** | **0.0002** |
| **AMOTL2** | **3:134070693-134094321** | **44.2539** | **20.0008** | **-1.14575** | **0.00035** |
| **CKB** | **14:103985995-103989448** | **52.3669** | **21.32** | **-1.29645** | **0.00035** |
| **FLNC** | **7:128469329-128550773** | **37.129** | **15.7097** | **-1.24089** | **0.00035** |
| **LMO2** | **11:33880121-33913836** | **3.0942** | **9.29468** | **1.58684** | **0.00035** |
| **DDX58** | **9:32454872-32526322** | **1.8323** | **4.83987** | **1.40132** | **0.0004** |
| **DUSP4** | **8:29190520-29208185** | **6.27976** | **13.8096** | **1.1369** | **0.0004** |
| **HSPB1** | **7:75931860-75933612** | **110.288** | **50.8884** | **-1.11586** | **0.00045** |
| **LIN7C** | **11:27516122-27528320** | **15.9151** | **7.58842** | **-1.06852** | **0.00045** |
| **COMP** | **19:18893582-18902123** | **8.88322** | **3.40808** | **-1.38212** | **0.0005** |
| **ZBTB7C** | **18:45553043-45937123** | **0.149535** | **1.01414** | **2.76171** | **0.00055** |
| **DACT1** | **14:59100684-59115039** | **21.1273** | **10.379** | **-1.02545** | **0.00065** |
| **MALL** | **2:110841446-110874143** | **6.92989** | **14.5298** | **1.06811** | **0.00065** |
| **CCDC136** | **7:128430810-128462186** | **5.36102** | **1.19772** | **-2.16222** | **0.0007** |
| **THBS2** | **6:169613348-169654139** | **67.6066** | **32.5462** | **-1.05467** | **0.0008** |
| **TNNT1** | **19:55644161-55660722** | **1.02306** | **0.0598316** | **-4.09584** | **0.0008** |
| **CNN1** | **19:11649531-11661138** | **3.35137** | **0.811653** | **-2.04581** | **0.00085** |
| **CITED2** | **6:139693392-139695757** | **68.6031** | **34.0373** | **-1.01116** | **0.00095** |
| **EREG** | **4:75230859-75254468** | **0.754483** | **2.26395** | **1.58528** | **0.001** |
| **MX1** | **21:42792230-42831141** | **0.782425** | **4.21775** | **2.43045** | **0.00105** |
| **OASL** | **12:121458094-121477045** | **0.255187** | **5.5378** | **4.43969** | **0.00105** |
| **HYDIN** | **16:70841280-71264625** | **0.962337** | **0.176096** | **-2.45018** | **0.0011** |
| **LRP2** | **2:169983618-170219195** | **0** | **0.540055** | **inf** | **0.0011** |
| **MT1E** | **16:56659386-56661024** | **133.205** | **274.099** | **1.04105** | **0.00115** |
| **NPTX1** | **17:78440639-78451643** | **2.06235** | **4.8782** | **1.24206** | **0.0013** |
| **HLF** | **17:53342372-53402426** | **1.57862** | **0.152429** | **-3.37245** | **0.0015** |
| **ACAA1** | **3:38080695-38178733** | **11.2895** | **4.59533** | **-1.29675** | **0.00155** |
| **ADRA2C** | **4:3768074-3770251** | **11.5676** | **5.39381** | **-1.10072** | **0.00175** |
| **ACTA2** | **10:90639490-90776816** | **127.449** | **55.27** | **-1.20535** | **0.0018** |
| **CAMK2B** | **7:44256748-44374176** | **0.133358** | **0.777907** | **2.5443** | **0.0018** |
| **HSPB7** | **1:16340465-16360545** | **76.7897** | **40.0746** | **-0.938225** | **0.0018** |
| **LOXL4** | **10:100007446-100028012** | **84.1985** | **166.736** | **0.985695** | **0.00185** |
| **SPTLC3** | **20:12989608-13151398** | **3.24986** | **8.95857** | **1.46289** | **0.0019** |
| **AIM1** | **6:106959729-107018472** | **0.462314** | **1.38423** | **1.58214** | **0.002** |
| **IGF2BP3** | **7:23338357-23510086** | **1.24434** | **6.08642** | **2.29022** | **0.002** |
| **CYR61** | **1:86046443-86050669** | **43.1341** | **22.5969** | **-0.932705** | **0.00205** |
| **PAPPA** | **9:118916052-119164601** | **20.7715** | **8.34183** | **-1.31617** | **0.00205** |
| **PLA2G4A** | **1:186798084-186958113** | **2.63129** | **6.25703** | **1.24971** | **0.00205** |
| **HERC5** | **4:89378267-89427314** | **0.120274** | **1.21947** | **3.34186** | **0.0021** |
| **ADCY8** | **8:131792546-132054672** | **0.308452** | **1.30445** | **2.08032** | **0.00215** |
| **NUP62CL** | **X:106366656-106449670** | **0** | **0.587258** | **inf** | **0.00215** |
| **ACTC1** | **15:35047284-35105124** | **7.38277** | **1.8493** | **-1.99719** | **0.0022** |
| **KCND3** | **1:112313283-112531777** | **0.882576** | **0.25651** | **-1.7827** | **0.00225** |
| **DSP** | **6:7541807-7586950** | **2.26003** | **1.09302** | **-1.04802** | **0.0024** |
| **UBE2E3** | **2:181831974-182264286** | **21.1877** | **45.9043** | **1.1154** | **0.00245** |
| **VAV3** | **1:108113781-108537229** | **0.0515539** | **0.68934** | **3.74106** | **0.00245** |
| **CRYAB** | **11:111779288-111797766** | **147.114** | **74.4264** | **-0.983051** | **0.0025** |
| **SH2B3** | **12:111843689-111889427** | **9.91015** | **18.3494** | **0.888756** | **0.00255** |
| **NRG1** | **8:31496901-32627154** | **16.2518** | **7.29204** | **-1.1562** | **0.0026** |
| **PELI2** | **14:56584531-56768244** | **2.16672** | **0.417653** | **-2.37513** | **0.0028** |
| **DNER** | **2:230222135-230579274** | **1.12754** | **2.95709** | **1.391** | **0.0029** |
| **GPR68** | **14:91698875-91720269** | **6.81219** | **13.4569** | **0.982155** | **0.00295** |
| **HMGA1** | **6:34204649-34214008** | **37.7381** | **69.55** | **0.882028** | **0.00295** |
| **AKR1B10** | **7:134200812-134226160** | **2.83409** | **7.02746** | **1.31012** | **0.00305** |
| **PRPS1** | **X:106871698-106894256** | **99.7844** | **50.9748** | **-0.96903** | **0.0031** |
| **GNG11** | **7:93551010-93557922** | **11.2282** | **21.1583** | **0.914098** | **0.00315** |
| **ZYX** | **7:143078172-143220542** | **18.4916** | **9.28806** | **-0.993424** | **0.0037** |
| **NDP** | **X:43808021-43832750** | **0.373239** | **2.29039** | **2.61742** | **0.0041** |
| **TMEM107** | **17:8076203-8079717** | **3718.53** | **7676.79** | **1.04577** | **0.00415** |
| **CTGF** | **6:132269315-132398533** | **117.392** | **64.8625** | **-0.855877** | **0.00435** |
| **ADAMTS5** | **21:28261694-28341390** | **10.9107** | **6.00952** | **-0.860424** | **0.0044** |
| **IRX3** | **16:54317215-54320710** | **17.3835** | **9.01455** | **-0.947391** | **0.0044** |
| **GDF15** | **19:18485540-18499987** | **31.2206** | **56.2992** | **0.850616** | **0.00475** |
| **TG** | **8:133879202-134147147** | **0.705459** | **0.0827603** | **-3.09155** | **0.00485** |
| **MCF2L** | **13:113548691-113754053** | **1.8877** | **0.47402** | **-1.99361** | **0.00515** |
| **FAM183A** | **1:43610823-43622067** | **0** | **0.761124** | **inf** | **0.0056** |
| **SAMD9** | **7:92728828-92747336** | **7.36512** | **13.4544** | **0.869299** | **0.0056** |
| **PTGFR** | **1:78695282-79006041** | **3.05254** | **6.3441** | **1.0554** | **0.00565** |
| **TFAP4** | **16:4307186-4323076** | **3.06056** | **0.767245** | **-1.99604** | **0.00565** |
| **HPGD** | **4:175411327-175444305** | **0.0156161** | **0.523427** | **5.06688** | **0.0057** |
| **FOXQ1** | **6:1312674-1314992** | **0.434921** | **1.8164** | **2.06226** | **0.0058** |
| **CDCP1** | **3:45123769-45187914** | **5.10175** | **9.49335** | **0.895926** | **0.006** |
| **TPM2** | **9:35681988-35691017** | **425.528** | **230.684** | **-0.88334** | **0.00605** |
| **STRA6** | **15:74471806-74628813** | **0.558766** | **1.86473** | **1.73866** | **0.0062** |
| **LBH** | **2:30454396-30546596** | **64.3002** | **35.8313** | **-0.843603** | **0.0064** |
| **RGS2** | **1:192778168-192781403** | **0.675541** | **3.37239** | **2.31966** | **0.00645** |
| **SPRY2** | **13:80910110-80915086** | **7.9049** | **15.2178** | **0.944936** | **0.00655** |
| **GJB2** | **13:20761608-20767037** | **0.684142** | **2.26593** | **1.72774** | **0.00715** |
| **MLPH** | **2:238394070-238463961** | **1.74749** | **4.29233** | **1.29647** | **0.00715** |
| **CXCL6** | **4:74702213-74714781** | **0.915695** | **3.09207** | **1.75564** | **0.00745** |
| **PPFIA2** | **12:81652044-82153332** | **0.817209** | **0.128825** | **-2.66529** | **0.00745** |
| **ACTG1** | **17:79476994-79494802** | **810.524** | **362.339** | **-1.16152** | **0.0075** |
| **IFIT3** | **10:90973325-91180758** | **6.1912** | **24.136** | **1.9629** | **0.0077** |
| **KIAA1199** | **15:81071683-81282219** | **409.863** | **199.716** | **-1.03719** | **0.00795** |
| **IL15RA** | **10:5990854-6020150** | **0.137821** | **0.672473** | **2.28668** | **0.0082** |
| **DCBLD2** | **3:98433173-98620533** | **42.809** | **77.7528** | **0.860979** | **0.00825** |
| **SLC2A1** | **1:43391051-43424530** | **17.87** | **8.75691** | **-1.02905** | **0.00825** |
| **IFI6** | **1:27992571-27998729** | **25.7545** | **46.0547** | **0.83852** | **0.00865** |
| **IFI44L** | **1:79085606-79111830** | **0.251288** | **1.89067** | **2.91148** | **0.0087** |
| **HRCT1** | **9:35904884-35907885** | **4.10413** | **1.37985** | **-1.57256** | **0.00885** |
| **TGFBR1** | **9:101866319-101916567** | **28.26** | **14.1176** | **-1.00127** | **0.0091** |
| **PMEPA1** | **20:56223447-56286592** | **3.52284** | **1.31161** | **-1.4254** | **0.0092** |
| **DHRS3** | **1:12627937-12677737** | **38.8129** | **22.1551** | **-0.808897** | **0.00935** |
| **PTX3** | **3:156893011-157251498** | **716.356** | **386.549** | **-0.890023** | **0.0094** |
| **LPHN2** | **1:81771844-82458120** | **7.6322** | **15.3649** | **1.00946** | **0.00945** |
| **PVRL4** | **1:161040677-161059389** | **0.962354** | **2.71024** | **1.49378** | **0.00945** |
| **EIF4A2** | **3:186499568-186524847** | **21709** | **58386.1** | **1.42733** | **0.0095** |
| **SLC6A15** | **12:85253491-85307394** | **0.0547357** | **0.847015** | **3.95183** | **0.0097** |
| **DMKN** | **19:35988121-36004560** | **2.66882** | **1.20004** | **-1.15312** | **0.00995** |
| **DUSP5** | **10:112257595-112271321** | **1.45537** | **3.26636** | **1.16629** | **0.01025** |
| **CARD16,CARD17,CASP1** | **11:104896169-104972158** | **2.56564** | **6.64316** | **1.37255** | **0.0103** |
| **MT1M** | **16:56662970-56667898** | **8.48485** | **17.4575** | **1.04089** | **0.0103** |
| **MCAM** | **11:119179240-119192231** | **2.68466** | **1.15903** | **-1.21183** | **0.0104** |
| **RRAD** | **16:66955581-66959547** | **0.336569** | **2.63178** | **2.96707** | **0.01075** |
| **C6orf1** | **6:34214156-34217247** | **17.6708** | **32.7636** | **0.89073** | **0.01125** |
| **LMCD1** | **3:7994491-8653610** | **21.271** | **10.6977** | **-0.991587** | **0.01145** |
| **AMIGO2** | **12:47469489-47630443** | **5.70956** | **2.94538** | **-0.954923** | **0.0118** |
| **TFPI2** | **7:93220884-93540577** | **6.0141** | **14.6771** | **1.28715** | **0.01205** |
| **ADAMTS15** | **11:130318868-130346532** | **0.342204** | **0.92125** | **1.42873** | **0.01215** |
| **SAMD13** | **1:84764048-84863514** | **0.0388877** | **0.544819** | **3.80839** | **0.01255** |
| **ARL15** | **5:53179774-53606412** | **4.18947** | **0.640747** | **-2.70894** | **0.01285** |
| **C15orf41** | **15:36871811-37110660** | **3.02699** | **6.85227** | **1.1787** | **0.0129** |
| **MARC2** | **1:220921566-220958150** | **2.33802** | **0.803379** | **-1.54114** | **0.0135** |
| **MYH9** | **22:36677153-36784063** | **118.877** | **69.7346** | **-0.769524** | **0.01355** |
| **SLC20A1** | **2:113403433-113421413** | **30.9124** | **51.4635** | **0.735366** | **0.01405** |
| **ALDH3A1** | **17:19640994-19652256** | **1.74528** | **4.23939** | **1.2804** | **0.01445** |
| **OXTR** | **3:8661085-9005457** | **5.21342** | **1.48844** | **-1.80842** | **0.01485** |
| **FCRLA** | **1:161676761-161684142** | **0.512339** | **1.38983** | **1.43974** | **0.01525** |
| **SH3TC2** | **5:148303201-148442726** | **0.907971** | **2.51625** | **1.47055** | **0.01525** |
| **LAMC2** | **1:183155147-183214035** | **14.2478** | **23.1412** | **0.699731** | **0.016** |
| **AFF3** | **2:100162322-100759201** | **0.806194** | **0.12953** | **-2.63784** | **0.0161** |
| **B4GALT1** | **9:33104079-33179981** | **44.5192** | **27.3415** | **-0.703334** | **0.01655** |
| **FZD2** | **17:42634719-42638556** | **2.62598** | **1.13908** | **-1.20499** | **0.01675** |
| **PTGS1** | **9:125132823-125158119** | **7.71716** | **13.7339** | **0.831597** | **0.0169** |
| **TBC1D19** | **4:26578058-26767026** | **7.11022** | **13.6495** | **0.940883** | **0.0169** |
| **FDXR** | **17:72858618-72869156** | **10.8923** | **18.3489** | **0.752388** | **0.0172** |
| **MAP3K5** | **6:136878184-137114237** | **1.64285** | **3.38356** | **1.04234** | **0.01755** |
| **OSR1** | **2:19551245-19558414** | **21.5464** | **12.0284** | **-0.840999** | **0.01755** |
| **USP18** | **22:18632665-18660164** | **1.51977** | **3.28903** | **1.11381** | **0.0176** |
| **TFAM** | **10:60144781-60159553** | **9.93258** | **19.9205** | **1.00402** | **0.0178** |
| **PTPN12** | **7:77166591-77269388** | **7.78753** | **13.9419** | **0.840193** | **0.01835** |
| **SLN** | **11:107578103-107595135** | **0** | **0.576286** | **inf** | **0.01835** |
| **ADAMTS1** | **21:28208065-28217728** | **176.149** | **86.5308** | **-1.02551** | **0.01855** |
| **FRAS1** | **4:78978723-79465423** | **0.697735** | **0.21602** | **-1.69151** | **0.01895** |
| **RSAD2** | **2:6980700-7038370** | **0.149651** | **1.91518** | **3.67781** | **0.01895** |
| **BTG2** | **1:203274618-203278730** | **3.87667** | **7.16982** | **0.88712** | **0.0192** |
| **ABRACL** | **6:139349667-139364738** | **12.65** | **22.6173** | **0.838285** | **0.01925** |
| **COL11A1** | **1:103342022-103574052** | **2.97882** | **1.33334** | **-1.1597** | **0.01955** |
| **ABL1** | **9:133588409-133763156** | **21.1512** | **13.1155** | **-0.689467** | **0.01975** |
| **PHYHIP** | **8:22077221-22089854** | **0.162176** | **0.815009** | **2.32926** | **0.01975** |
| **SLCO4A1** | **20:61272070-61317137** | **0.378682** | **1.19385** | **1.65656** | **0.0198** |
| **CLK1** | **2:201717731-201729422** | **18.5627** | **31.3689** | **0.756931** | **0.0199** |
| **DMXL1** | **5:118342041-118585013** | **5.17669** | **10.5772** | **1.03086** | **0.02015** |
| **SLC26A4** | **7:107294420-107358254** | **0.0470358** | **0.501289** | **3.41381** | **0.02025** |
| **ARNT2** | **15:80696691-80890278** | **5.15485** | **2.67704** | **-0.945292** | **0.0203** |
| **NNMT** | **11:113930314-114227293** | **317.871** | **191.52** | **-0.730947** | **0.02065** |
| **ERCC2** | **19:45836691-45874176** | **14.6518** | **8.10585** | **-0.854042** | **0.02075** |
| **PFDN4** | **20:52824385-52844591** | **27.2258** | **48.1065** | **0.821259** | **0.02115** |
| **RCAN1** | **21:35885439-35987598** | **72.0043** | **45.0935** | **-0.675162** | **0.02115** |
| **ALDH7A1** | **5:125877532-125931110** | **14.5693** | **7.91725** | **-0.879862** | **0.02135** |
| **FAM171A2** | **17:42430582-42441243** | **1.3231** | **0.454179** | **-1.54258** | **0.02145** |
| **PTGIS** | **20:48120405-48184683** | **53.4972** | **29.8191** | **-0.843224** | **0.0215** |
| **FAM111B** | **11:58874657-58895131** | **3.45192** | **6.33043** | **0.874903** | **0.0217** |
| **SOWAHC** | **2:110371910-110376563** | **2.67569** | **1.43272** | **-0.901151** | **0.02185** |
| **TNFRSF10C** | **8:22941867-22979089** | **1.61776** | **3.34865** | **1.04958** | **0.02215** |
| **CYB561D1** | **1:110036289-110045554** | **3.11928** | **5.9997** | **0.943678** | **0.0223** |
| **LIMCH1** | **4:41361623-41702062** | **15.0383** | **7.57607** | **-0.989123** | **0.0224** |
| **AKAP6** | **14:32798478-33300567** | **1.51925** | **0.488455** | **-1.63706** | **0.0225** |
| **TMC4** | **19:54663783-54676944** | **1.48717** | **0.341986** | **-2.12056** | **0.02265** |
| **RRAGC** | **1:39303869-39325495** | **8.73034** | **14.6432** | **0.74612** | **0.02285** |
| **BRCA2** | **13:32889610-32974404** | **3.70547** | **6.97341** | **0.912208** | **0.02315** |
| **PMAIP1** | **18:57567179-57571538** | **3.27576** | **6.37832** | **0.961348** | **0.0232** |
| **MYC** | **8:128747679-128753674** | **16.1218** | **9.74559** | **-0.726188** | **0.0233** |
| **NDUFS4** | **5:52856462-52979169** | **54.7907** | **88.0887** | **0.685026** | **0.0236** |
| **NGF** | **1:115825654-115910693** | **13.8675** | **7.62928** | **-0.862089** | **0.02365** |
| **TFPI** | **2:187866602-188430487** | **20.5626** | **37.5689** | **0.869515** | **0.02375** |
| **SGK1** | **6:134490383-134639250** | **7.73947** | **13.7427** | **0.828359** | **0.02385** |
| **SSX2IP** | **1:85109389-85156486** | **5.32568** | **9.62503** | **0.853826** | **0.02395** |
| **FAM46B** | **1:27331510-27339327** | **3.0772** | **1.47952** | **-1.05649** | **0.024** |
| **MMP3** | **11:102617698-102714534** | **1.22667** | **4.74433** | **1.95146** | **0.024** |
| **ERRFI1** | **1:8064463-8086368** | **61.3811** | **98.1516** | **0.677217** | **0.0241** |
| **PPAP2B** | **1:56959309-57110974** | **201.008** | **117.599** | **-0.77338** | **0.02445** |
| **ABCA6** | **17:67074842-67138029** | **0.375663** | **1.32404** | **1.81744** | **0.0245** |
| **FBN2** | **5:127593551-128369335** | **16.0295** | **9.27804** | **-0.788837** | **0.02455** |
| **WISP2** | **20:43285091-43380409** | **131.834** | **81.9759** | **-0.68545** | **0.0246** |
| **ARHGAP18** | **6:129897024-130031370** | **5.64311** | **10.0776** | **0.836592** | **0.02465** |
| **SDPR** | **2:192699027-193060435** | **1.5469** | **2.98365** | **0.947703** | **0.0247** |
| **SCG2** | **2:224461657-224467221** | **1.57359** | **3.15604** | **1.00406** | **0.0247** |
| **ARRDC3** | **5:90664518-90679176** | **16.8094** | **30.1493** | **0.842861** | **0.02535** |
| **H2AFX** | **11:118964563-118966177** | **31.1716** | **19.18** | **-0.700628** | **0.02545** |
| **MSX2** | **5:174151535-174157896** | **1.868** | **3.99075** | **1.09517** | **0.0255** |
| **SEPHS2** | **16:30454951-30457502** | **3.38843** | **1.7351** | **-0.9656** | **0.02615** |
| **EFR3B** | **2:25264998-25379117** | **0.795313** | **1.68917** | **1.08672** | **0.0262** |
| **SLC4A4** | **4:72052494-72437804** | **6.72928** | **3.7638** | **-0.838263** | **0.02625** |
| **SLC1A5** | **19:47222763-47291851** | **115.23** | **67.2578** | **-0.776746** | **0.02635** |
| **CCDC124** | **19:18043824-18054800** | **9.36044** | **4.68326** | **-0.999063** | **0.0264** |
| **SLC13A5** | **17:6588031-6616886** | **0.656263** | **0.067414** | **-3.28316** | **0.0265** |
| **KLHL13** | **X:117031775-117251303** | **4.7294** | **2.40291** | **-0.976878** | **0.02655** |
| **SEMA3A** | **7:83585092-84122174** | **17.0378** | **35.9686** | **1.078** | **0.02665** |
| **PUS7L** | **12:44122272-44152620** | **6.05806** | **10.6102** | **0.808523** | **0.0271** |
| **ACTN1** | **14:69340845-69446157** | **54.4139** | **34.4372** | **-0.660007** | **0.0276** |
| **F2RL1** | **5:76114757-76131140** | **0.247821** | **0.890494** | **1.84531** | **0.0276** |
| **PRSS3** | **9:33750514-33920402** | **1.54883** | **9.16925** | **2.56562** | **0.0278** |
| **FSTL3** | **19:676380-683389** | **15.7295** | **9.1253** | **-0.785526** | **0.0279** |
| **LY96** | **8:74903586-74941322** | **25.3918** | **43.595** | **0.779804** | **0.02795** |
| **IL8** | **4:74606222-74609433** | **0.692347** | **1.92188** | **1.47295** | **0.028** |
| **WDHD1** | **14:55405667-55493823** | **6.47319** | **10.6126** | **0.713233** | **0.028** |
| **GRIP1** | **12:66741210-67197966** | **0.548415** | **0.0767292** | **-2.83742** | **0.0283** |
| **ACAN** | **15:89340643-89418585** | **50.6099** | **29.8087** | **-0.763683** | **0.0284** |
| **ISOC1** | **5:128430443-128449776** | **7.34335** | **4.12217** | **-0.833035** | **0.0284** |
| **PDE1C** | **7:31790792-32338941** | **28.8693** | **17.1452** | **-0.751731** | **0.0285** |
| **PLK5** | **19:1524072-1535455** | **0.905843** | **0** | **-inf** | **0.02865** |
| **CENPE** | **4:104026962-104119566** | **9.33248** | **14.9611** | **0.680885** | **0.02875** |
| **ID3** | **1:23884408-23886285** | **281.769** | **175.899** | **-0.679769** | **0.0288** |
| **PODXL** | **7:131185020-131242976** | **14.5966** | **22.7147** | **0.63799** | **0.02895** |
| **TMEM158** | **3:45265849-45267782** | **2.9622** | **5.5747** | **0.912225** | **0.02895** |
| **SLC25A23** | **19:6435992-6482568** | **5.24287** | **2.74609** | **-0.932979** | **0.02905** |
| **SPARCL1** | **4:88394486-88452213** | **0.0947454** | **0.577289** | **2.60717** | **0.0291** |
| **PRELP** | **1:203444955-203460480** | **1.63899** | **0.859568** | **-0.931126** | **0.0294** |
| **ZBTB18** | **1:244212198-244220778** | **1.77175** | **0.87639** | **-1.01553** | **0.02985** |
| **ACYP1** | **14:75519602-75545126** | **7.49433** | **20.2062** | **1.43093** | **0.0299** |
| **SLC27A3** | **1:153746829-153752633** | **4.08278** | **1.93642** | **-1.07616** | **0.02995** |
| **FGF2** | **4:123747862-123844123** | **64.0813** | **38.8809** | **-0.720843** | **0.03015** |
| **BZRAP1** | **17:56378591-56494956** | **0.530395** | **0.0282976** | **-4.22832** | **0.0305** |
| **ANXA10** | **4:168867824-169108841** | **0.240857** | **1.72843** | **2.84322** | **0.03055** |
| **WDR63** | **1:85464829-85598821** | **0.246107** | **0.936623** | **1.92818** | **0.03065** |
| **KRT19** | **17:39679867-39684560** | **3.40123** | **6.38562** | **0.908772** | **0.0307** |
| **NFATC4** | **14:24834878-24848810** | **61.9727** | **29.452** | **-1.07327** | **0.03075** |
| **HSPA5** | **9:127997131-128003609** | **69.968** | **44.362** | **-0.657372** | **0.0311** |
| **RCAN2** | **6:46188474-46459709** | **0.118388** | **0.791957** | **2.74189** | **0.03115** |
| **TINAGL1** | **1:32042115-32053288** | **3.7865** | **2.10887** | **-0.844393** | **0.0312** |
| **SUGCT** | **7:40174574-40900362** | **3.24835** | **7.16124** | **1.1405** | **0.03125** |
| **C12orf56** | **12:64616116-64790845** | **0.228072** | **0.767882** | **1.75139** | **0.0313** |
| **AMIGO1** | **1:110046796-110052360** | **1.87239** | **0.973828** | **-0.943139** | **0.03135** |
| **RGS4** | **1:163038564-163046592** | **1.16853** | **3.52934** | **1.59471** | **0.0314** |
| **ADM** | **11:10326226-10328945** | **115.778** | **71.3269** | **-0.698843** | **0.03155** |
| **EGF** | **4:110834039-110933422** | **1.28144** | **3.54178** | **1.46671** | **0.0318** |
| **ZNF43** | **19:21987751-22034927** | **2.27849** | **4.79602** | **1.07376** | **0.03185** |
| **PXDC1** | **6:3722847-3754105** | **79.123** | **49.1735** | **-0.686217** | **0.0322** |
| **DUSP1** | **5:172185228-172204777** | **45.1107** | **28.9646** | **-0.639179** | **0.03225** |
| **VARS** | **6:31745231-31763730** | **12.279** | **7.63287** | **-0.685901** | **0.03235** |
| **SNAI2** | **8:49830248-49834860** | **14.7232** | **9.11317** | **-0.692071** | **0.03245** |
| **ATOH8** | **2:85978466-86015189** | **41.5505** | **26.6495** | **-0.64076** | **0.03255** |
| **PPFIA4** | **1:202995625-203047868** | **4.78673** | **2.64767** | **-0.854316** | **0.0331** |
| **GMNN** | **6:24775158-24786327** | **11.7899** | **19.9747** | **0.760619** | **0.0342** |
| **C3** | **19:6677714-6737614** | **0.186319** | **0.787687** | **2.07985** | **0.0345** |
| **CHIC1** | **X:72782983-72906937** | **2.41415** | **0.94695** | **-1.35016** | **0.0345** |
| **FAM96A** | **15:64364757-64386217** | **32.3771** | **19.3257** | **-0.744452** | **0.0346** |
| **PAG1** | **8:81880044-82024335** | **1.06537** | **3.02788** | **1.50695** | **0.0346** |
| **CALD1** | **7:134429002-134655479** | **265.796** | **158.071** | **-0.749748** | **0.0347** |
| **PITPNC1** | **17:65373574-65693372** | **2.46039** | **5.15246** | **1.06638** | **0.035** |
| **MFSD3** | **8:145734456-145736596** | **9.57489** | **5.38058** | **-0.831494** | **0.03535** |
| **C18orf54** | **18:51850727-51911588** | **1.14035** | **2.59207** | **1.18463** | **0.0355** |
| **DENND1A** | **9:126118448-126692431** | **2.60624** | **5.26085** | **1.01333** | **0.03575** |
| **C1orf198** | **1:230972696-231005335** | **19.5195** | **11.3126** | **-0.786983** | **0.03605** |
| **GAS6** | **13:114518602-114569806** | **127.702** | **82.4004** | **-0.632052** | **0.03615** |
| **SAT1** | **X:23801274-23804343** | **44.864** | **69.4187** | **0.629766** | **0.03615** |
| **CLEC2B** | **12:10005582-10022735** | **0.405155** | **1.33736** | **1.72285** | **0.03625** |
| **SPTSSA** | **14:34393436-34931980** | **2.2229** | **1.12414** | **-0.983625** | **0.03625** |
| **FAM154B** | **15:82555150-82577271** | **1.11016** | **0.0366511** | **-4.92076** | **0.03715** |
| **NOP56** | **20:2632018-2644865** | **377.413** | **2240.13** | **2.56936** | **0.0373** |
| **ABCC1** | **16:16043433-16236931** | **11.6303** | **7.15628** | **-0.700612** | **0.0374** |
| **CD36** | **7:79959507-80308593** | **1.80954** | **3.99847** | **1.14382** | **0.03765** |
| **RPL3** | **22:39708886-39716394** | **397.334** | **256.424** | **-0.631823** | **0.0379** |
| **ZDHHC9** | **X:128937184-128977885** | **13.3714** | **8.24645** | **-0.697301** | **0.03835** |
| **CCDC138** | **2:109403132-109501933** | **1.7024** | **3.36887** | **0.98469** | **0.03865** |
| **SEC14L1** | **17:75082797-75213261** | **1572.65** | **2644.53** | **0.749808** | **0.03895** |
| **TOPBP1** | **3:133317018-133380821** | **9.93745** | **17.269** | **0.797234** | **0.039** |
| **IL7R** | **5:35852796-35879706** | **2.62841** | **5.10073** | **0.956514** | **0.0391** |
| **IFIT1** | **10:90973325-91180758** | **6.44325** | **19.3393** | **1.58568** | **0.03955** |
| **ZER1** | **9:131486723-131534693** | **8.16763** | **3.89584** | **-1.06798** | **0.03955** |
| **GADD45B** | **19:2476119-2478877** | **40.2904** | **25.657** | **-0.651085** | **0.03985** |
| **RIIAD1** | **1:151673501-151702281** | **0.0556836** | **1.14643** | **4.36375** | **0.04015** |
| **BLOC1S2** | **10:102033712-102046575** | **30.8592** | **48.1438** | **0.641649** | **0.0402** |
| **CTPS1** | **1:41445006-41478466** | **9.99192** | **5.33484** | **-0.905317** | **0.0403** |
| **GNPDA1** | **5:141371313-141392606** | **11.5489** | **18.4802** | **0.678226** | **0.0403** |
| **MBOAT7** | **19:54677105-54693733** | **16.4952** | **9.00335** | **-0.873514** | **0.0403** |
| **TGFBR2** | **3:30647993-30737488** | **27.6254** | **42.0703** | **0.606807** | **0.0404** |
| **SCUBE3** | **6:35181700-35221295** | **23.0164** | **15.1535** | **-0.603018** | **0.04045** |
| **TTC25** | **17:40086887-40117648** | **1.4862** | **0.372298** | **-1.9971** | **0.04045** |
| **PYGM** | **11:64513860-64527769** | **0.787732** | **0.0717692** | **-3.45627** | **0.04065** |
| **FHL3** | **1:38462400-38471278** | **3.6226** | **1.83723** | **-0.979495** | **0.04075** |
| **BRSK1** | **19:55793439-55823901** | **1.50317** | **0.53511** | **-1.49011** | **0.04085** |
| **FBLN5** | **14:92335755-92414345** | **6.76313** | **4.06041** | **-0.736066** | **0.04095** |
| **SLC43A2** | **17:1472560-1533052** | **4.51049** | **2.3296** | **-0.9532** | **0.04145** |
| **MRPL39** | **21:26957967-26979829** | **16.0056** | **25.7324** | **0.685009** | **0.04175** |
| **ATP2A2** | **12:110718560-110789058** | **49.3019** | **31.8498** | **-0.630361** | **0.04185** |
| **CMKLR1** | **12:108681820-108733189** | **16.3552** | **26.3871** | **0.690082** | **0.0419** |
| **NDST1** | **5:149855093-149937773** | **23.1026** | **14.2548** | **-0.696611** | **0.0421** |
| **GBP3** | **1:89472348-89488577** | **9.16869** | **15.2706** | **0.735973** | **0.0424** |
| **HERPUD1** | **16:56965959-56977798** | **41.92** | **26.6547** | **-0.653248** | **0.0432** |
| **FAM19A5** | **22:48885271-49246724** | **5.28393** | **2.84888** | **-0.891216** | **0.04345** |
| **NAA30** | **14:57857261-57882635** | **4.17494** | **7.12668** | **0.771475** | **0.04385** |
| **SAMD4A** | **14:55032690-55260033** | **9.36105** | **5.67004** | **-0.723313** | **0.04405** |
| **CCDC82** | **11:96085932-96123087** | **4.32021** | **7.60542** | **0.815925** | **0.04415** |
| **COL14A1** | **8:121072018-121384759** | **13.1691** | **19.8678** | **0.593279** | **0.04415** |
| **TLCD2** | **17:1610185-1614203** | **2.46** | **0.956779** | **-1.3624** | **0.04425** |
| **DOCK8** | **9:213107-746105** | **0.154208** | **0.766756** | **2.31389** | **0.04435** |
| **ERICH1** | **8:564745-1212580** | **4.08637** | **2.01603** | **-1.0193** | **0.0445** |
| **SLC1A1** | **9:4490443-4666674** | **8.67299** | **13.4362** | **0.631528** | **0.0446** |
| **MAPKAPK3** | **3:50643920-50686720** | **40.2369** | **62.9049** | **0.644653** | **0.04465** |
| **MT1F** | **16:56691605-56694610** | **0.412703** | **1.73845** | **2.07462** | **0.04465** |
| **RARRES3** | **11:63304280-63313934** | **2.27186** | **5.23408** | **1.20406** | **0.0447** |
| **WDR75** | **2:190306158-190340291** | **10.44** | **17.3071** | **0.729242** | **0.0449** |
| **ARAP2** | **4:35949842-36275842** | **0.0739184** | **0.678952** | **3.1993** | **0.0453** |
| **TPM1** | **15:63334452-63364114** | **654.38** | **408.219** | **-0.680786** | **0.04565** |
| **MSRB1** | **16:1987860-1993327** | **6.62393** | **3.11722** | **-1.08743** | **0.0459** |
| **CCDC85B** | **11:65657176-65668266** | **68.3761** | **37.2698** | **-0.875486** | **0.0462** |
| **IGFBP2** | **2:217497550-217529159** | **12.1466** | **19.738** | **0.700424** | **0.04625** |
| **MANBAL** | **20:35918040-35945663** | **8.89899** | **5.19496** | **-0.776528** | **0.04645** |
| **ZNF281** | **1:200374067-200452680** | **17.2311** | **11.1822** | **-0.623813** | **0.04655** |
| **GPR153** | **1:6307405-6321035** | **3.38856** | **5.34823** | **0.658387** | **0.0479** |
| **ZNF445** | **3:44481261-44519162** | **3.10556** | **1.71648** | **-0.855395** | **0.0479** |
| **GYS1** | **19:49471381-49496567** | **21.9232** | **13.1997** | **-0.731952** | **0.04795** |
| **CENPK** | **5:64813592-64858998** | **18.7779** | **30.1378** | **0.682539** | **0.04815** |
| **ELOVL5** | **6:53132195-53213947** | **49.0561** | **30.7841** | **-0.672249** | **0.04845** |
| **ETV4** | **17:41561232-41687706** | **0.203021** | **0.866459** | **2.09351** | **0.0486** |
| **SELM** | **22:31500757-31516055** | **87.1065** | **57.4677** | **-0.60003** | **0.0488** |
| **CD58** | **1:117057156-117113661** | **5.82328** | **10.2645** | **0.817761** | **0.04905** |
| **MDM2** | **12:69201841-69365350** | **48.866** | **97.6323** | **0.998528** | **0.0492** |
| **DDAH1** | **1:85731930-86043933** | **88.5534** | **52.865** | **-0.744235** | **0.0495** |
| **RING1** | **6:33176065-33180800** | **11.7726** | **7.32445** | **-0.684639** | **0.04955** |
| **NCL** | **2:232318241-232348352** | **57.048** | **35.1595** | **-0.698261** | **0.0501** |
| **STX17** | **9:102668914-102737639** | **4.64354** | **7.80878** | **0.749872** | **0.05035** |
| **DEPTOR** | **8:120879658-121063152** | **1.31821** | **0.533724** | **-1.30441** | **0.0505** |
| **NCOA7** | **6:126102279-126252266** | **8.01145** | **15.0739** | **0.911918** | **0.0507** |
| **STAT4** | **2:191886251-192016322** | **3.25801** | **1.83899** | **-0.825079** | **0.0511** |
| **PDLIM1** | **10:96997328-97050781** | **2.61665** | **1.21878** | **-1.10228** | **0.0512** |
| **LMOD1** | **1:201857218-201915715** | **1.44729** | **0.400778** | **-1.85248** | **0.05135** |
| **SASH1** | **6:148558720-148873187** | **10.5857** | **7.10393** | **-0.575428** | **0.05165** |
| **MSC** | **8:72740401-73034094** | **9.75188** | **5.19818** | **-0.907674** | **0.0518** |
| **SLC35D2** | **9:99082986-99145992** | **6.35598** | **10.7608** | **0.759599** | **0.0518** |
| **CCDC102B** | **18:66340924-66722426** | **0.140584** | **0.6611** | **2.23344** | **0.05185** |
| **MRPS28,TPD52** | **8:80830951-81143467** | **33.2583** | **53.4593** | **0.684727** | **0.0521** |
| **SNX2** | **5:122110690-122165925** | **22.8122** | **34.4647** | **0.595316** | **0.0521** |
