## Supplementary table 4 for "Dissecting *ARL15* Function in Rheumatoid Arthritis: Insights from *Ex Vivo* and *In Vitro* Synovial Fibroblast Models"

| **Supplementary table 4: 87 differentially regulated genes (P<0.05) obtained upon *ARL15* knockdown in synovial fluid derived OASF1** | | | | | | |
| --- | --- | --- | --- | --- | --- | --- |
| **Gene** | **locus** | **OASF1**  **Control FPKM** | | **OASF1**  **(ARL15_siRNA)FPKM** | **log2(fold change)** | **P_value_** |
| **AGR2** | **7:16831434-16873057** | **0** | | **0.307257** | **inf** | **0.00005** |
| **CSN2** | **4:70820973-70831434** | **0** | | **1.74251** | **inf** | **0.00005** |
| **FHIT** | **3:59735035-61237133** | **0.458881** | | **0** | **-inf** | **0.00005** |
| **LALBA** | **12:48961466-48963849** | **0** | | **1.84435** | **inf** | **0.00005** |
| **NPIPA5** | **16:15457515-15474904** | **0.935012** | | **8.42124** | **3.17098** | **0.00005** |
| **PPBP** | **4:74852754-74853914** | **0** | | **0.429009** | **inf** | **0.00005** |
| **ARL15** | **5:53179774-53606415** | **3.82141** | | **0.957777** | **-1.99634** | **0.0002** |
| **TEKT3** | **17:15207127-15244958** | **0** | | **0.355834** | **inf** | **0.00035** |
| **FAM96A** | **15:64364757-64386217** | **19.4223** | | **11.2181** | **-0.791886** | **0.0004** |
| **NPTX1** | **17:78440723-78451643** | **0.702237** | | **1.44524** | **1.04127** | **0.0005** |
| **MX1** | **21:42792230-42831141** | **10.2105** | | **5.31932** | **-0.940738** | **0.0007** |
| **TFF3** | **21:43731776-43735761** | **0** | | **0.364808** | **inf** | **0.00085** |
| **LST1** | **6:31553900-31560762** | **0.0257638** | | **0.246991** | **3.26104** | **0.0013** |
| **VCAM1** | **1:101185297-101204601** | **2.44492** | | **4.10119** | **0.746258** | **0.0013** |
| **SKAP2** | **7:26706680-27034858** | **5.96948** | | **3.002** | **-0.991681** | **0.00195** |
| **PHGR1** | **15:40643233-40648635** | **0** | | **0.433178** | **inf** | **0.00215** |
| **CSN3** | **4:71108304-71117145** | **0** | | **0.246459** | **inf** | **0.00275** |
| **CCDC171** | **9:15552894-16061661** | **0.413305** | | **1.00315** | **1.27926** | **0.0035** |
| **ANKRD9** | **14:102969822-102976136** | **4.72267** | | **8.01586** | **0.763256** | **0.00355** |
| **CYP2D6** | **22:42481528-42540576** | **0.0643072** | | **1.48103** | **4.52548** | **0.0048** |
| **NPIPA2** | **16:14841922-14859270** | **0.193312** | | **1.49317** | **2.94937** | **0.0058** |
| **SIDT1** | **3:113251142-113348425** | **0.397852** | | **0.0146782** | **-4.76048** | **0.0063** |
| **ESYT3** | **3:138153427-138200528** | **0.684315** | | **0.255737** | **-1.42** | **0.0082** |
| **GHR** | **5:42423878-42721979** | **0.737827** | | **1.33648** | **0.857084** | **0.0094** |
| **PRLR** | **5:35048860-35230794** | **0.14959** | | **0.685763** | **2.19669** | **0.0096** |
| **C18orf54** | **18:51850727-51911588** | **1.99102** | | **0.879491** | **-1.17876** | **0.0107** |
| **UBE2U** | **1:64669309-64733051** | **0.229845** | | **0** | **-inf** | **0.0115** |
| **ADH1B** | **4:100226120-100242558** | **12.7887** | | **19.328** | **0.595817** | **0.01405** |
| **NPIPB15** | **16:74411775-74426019** | **5.70058** | | **3.45248** | **-0.723479** | **0.01435** |
| **C8orf34** | **8:69215702-69731257** | **0.529541** | | **1.2079** | **1.18968** | **0.01855** |
| **GBP1** | **1:89518001-89531043** | **14.2296** | | **8.92517** | **-0.67294** | **0.01965** |
| **C9orf92** | **9:16203932-16276311** | **0.32899** | | **0** | **-inf** | **0.02035** |
| **ZNF107** | **7:64126510-64171404** | **0.678765** | | **0.421675** | **-0.68678** | **0.02245** |
| **LGALS4** | **19:39292310-39304004** | **0.15173** | | **1.24073** | **3.03161** | **0.02275** |
| **TRH** | **3:129693147-129696781** | **3.27848** | | **4.82151** | **0.556458** | **0.02285** |
| **STYK1** | **12:10771537-10826917** | **0.781831** | | **0.058928** | **-3.72983** | **0.02305** |
| **ROPN1L** | **5:10441401-10472141** | **0.657117** | | **0.14523** | **-2.17781** | **0.02325** |
| **PIEZO2** | **18:10661929-11148935** | **0.320304** | | **0.774971** | **1.2747** | **0.0241** |
| **ATL2** | **2:38521178-38604514** | **6.55132** | | **9.84929** | **0.588235** | **0.03015** |
| **GAS2L3** | **12:100967460-101022064** | **2.94362** | | **1.76552** | **-0.737497** | **0.0315** |
| **TMEM121** | **14:105992939-105996539** | **0.691073** | | **1.36212** | **0.978944** | **0.03185** |
| **MACC1** | **7:19958603-20257027** | **0.0766622** | | **0.412537** | **2.42794** | **0.032** |
| **C9orf41** | **9:77567883-77643339** | **3.55939** | | **2.09831** | **-0.762402** | **0.03335** |
| **HIST1H2BJ** | **6:27091862-27100562** | **1.41727** | | **2.97052** | **1.0676** | **0.03415** |
| **BAIAP2L2** | **22:38480895-38506677** | **0.828233** | | **1.66244** | **1.00519** | **0.03455** |
| **RASGRP3** | **2:33661390-33789817** | **0.0919441** | | **0.328342** | **1.83637** | **0.0352** |
| **ATP1A3** | **19:42470733-42501649** | **1.02339** | | **0.536667** | **-0.931252** | **0.0362** |
| **CRYM** | **16:21244985-21329912** | **0.0689312** | | **0.388171** | **2.49347** | **0.0372** |
| **CD83** | **6:14103603-14137149** | **0.344635** | | **0.640005** | **0.893014** | **0.03735** |
| **NAPRT1** | **8:144655659-144660819** | **1.3251** | | **2.34864** | **0.825724** | **0.03785** |
| **MMP1** | **11:102617698-102714534** | **15.1935** | | **20.4253** | **0.426903** | **0.03845** |
| **LHFPL4** | **3:9543480-9595486** | **0.307104** | | **0** | **-inf** | **0.04** |
| **NOS3** | **7:150688082-150721586** | **0.137961** | | **0.60491** | **2.13246** | **0.04095** |
| **GRASP** | **12:52400723-52409673** | **0.341207** | | **0.0806634** | **-2.08066** | **0.0412** |
| **CCNG1** | **5:162864574-162887146** | **65.6434** | | **94.5087** | **0.525796** | **0.04245** |
| **SWSAP1** | **19:11478798-11487627** | **0.438211** | | **0.842515** | **0.943078** | **0.04275** |
| **ACTG2** | **2:74119440-74146992** | **0.452386** | | **0.151836** | **-1.57504** | **0.04305** |
| **NRG2** | **5:139226363-139422884** | **0.157177** | | **0.389748** | **1.31015** | **0.04365** |
| **ATG9B** | **7:150688082-150721586** | **0.41958** | | **0.106933** | **-1.97224** | **0.04375** |
| **IGFBP2** | **2:217497550-217529159** | **0.616866** | | **1.20724** | **0.968689** | **0.04375** |
| **LGI2** | **4:25000468-25032501** | **0.568062** | | **0.26918** | **-1.07748** | **0.04375** |
| **CA4** | **17:58227296-58248260** | **0.130202** | | **0.413896** | **1.66852** | **0.044** |
| **NAT14,ZNF628** | **19:55986363-55998939** | **7.94408** | | **11.9464** | **0.588628** | **0.044** |
| **FGFR4** | **5:176513886-176525145** | **0.273276** | | **0.0603857** | **-2.17808** | **0.0441** |
| **ZNF28,ZNF468** | **19:53300661-53360902** | **4.02588** | | **6.04961** | **0.587537** | **0.04445** |
| **STOX2** | **4:184774583-184944679** | **0.123955** | | **0.484736** | **1.96739** | **0.045** |
| **MEIOB** | **16:1845620-1934295** | **0.0140374** | | **0.435666** | **4.95588** | **0.04605** |
| **CYFIP2** | **5:156687131-157002783** | **1.39496** | | **2.11932** | **0.603373** | **0.04615** |
| **C4orf48** | **4:2043688-2045697** | **13.7571** | | **19.7269** | **0.519987** | **0.04795** |
| **RASAL3** | **19:15562434-15575382** | **0.0412452** | | **0.299226** | **2.85894** | **0.05** |
| **ZNF608** | **5:123972607-124084500** | **2.60348** | | **3.78367** | **0.539346** | **0.05035** |
| **NCF1** | **7:74071993-74306731** | **0.0549441** | | **0.275851** | **2.32785** | **0.0508** |
| **ANKRD30B** | **18:14728270-14854033** | **0.174769** | | **0.434628** | **1.31433** | **0.0509** |
| **CDH2** | **18:25530929-25757410** | **6.30574** | | **8.91819** | **0.500085** | **0.0512** |
| **ZDHHC21** | **9:14611068-14693469** | **2.37335** | | **1.39304** | **-0.768682** | **0.0512** |
| **MAN1C1** | **1:25943958-26112698** | **2.30284** | | **3.39883** | **0.561626** | **0.05125** |
| **PRSS36** | **16:31150245-31161415** | **1.33011** | | **0.619497** | **-1.10238** | **0.0517** |
| **DUSP15** | **20:30435439-30539895** | **0.346255** | | **0.0724615** | **-2.25655** | **0.0519** |
| **C5AR1** | **19:47793279-47825323** | **0.170143** | | **0.491274** | **1.52978** | **0.0528** |
| **KRT8** | **12:53290976-53346686** | **1.76034** | | **3.69271** | **1.06883** | **0.05295** |
| **HMGA2** | **12:66149962-66360075** | **6.99401** | | **4.22343** | **-0.727706** | **0.05325** |
| **RLTPR** | **16:67678821-67694713** | **0.0841091** | | **0.441844** | **2.3932** | **0.0535** |
| **LRRCC1** | **8:86019381-86058311** | **2.06663** | | **1.47077** | **-0.490705** | **0.05425** |
| **BTBD3** | **20:11871370-11907257** | **2.86731** | | **4.58378** | **0.67684** | **0.0545** |
| **ZNF418** | **19:58433251-58446761** | **1.87467** | | **2.8893** | **0.624081** | **0.0549** |
| **RASGRP2** | **11:64494382-64512928** | **0.540583** | | **0.177339** | **-1.60801** | **0.05535** |
| **HLA-F** | **6:29690551-29716826** | **5.84131** | | **8.53867** | **0.547718** | **0.0555** |
