## Supplementary table 5 for "Dissecting *ARL15* Function in Rheumatoid Arthritis: Insights from *Ex Vivo* and *In Vitro* Synovial Fibroblast Models"

**Supplementary table 5:** 56 differentially expressed genes upon *ARL15* KD in MH7A cells

| **S.No** | **Ensemble ID** | **Gene** | **log2 FC** | **P-adj. value** |
| --- | --- | --- | --- | --- |
| **Downregulated genes** | | | | |
| 1 | ENSG00000185305 | *ARL15* | -3.9 | 1.39E-49 |
| 2 | ENSG00000147065 | *MSN* | -1.2 | 4.05E-23 |
| 3 | ENSG00000149547 | *EI24* | -1.2 | 1.67E-15 |
| 4 | ENSG00000003436 | *TFPI* | -1.3 | 3.42E-15 |
| 5 | ENSG00000170348 | *TMED10* | -1.2 | 3.68E-15 |
| 6 | ENSG00000166881 | *TMEM194* (*NEMP1*) | -1.1 | 8.35E-15 |
| 7 | ENSG00000106538 | *RARRES2* | -1.4 | 7.81E-14 |
| 8 | ENSG00000173706 | *HEG1* | -1.1 | 2.82E-12 |
| 9 | ENSG00000100304 | *TTLL12* | -1.0 | 3.94E-10 |
| 10 | ENSG00000128510 | *CPA4* | -1.0 | 1.72E-09 |
| 11 | ENSG00000214049 | *UCA1* | -1.2 | 2.16E-08 |
| 12 | ENSG00000085117 | *CD82* | -1.0 | 1.45E-07 |
| 13 | ENSG00000136869 | *TLR4* | -1.0 | 1.30E-06 |
| 14 | ENSG00000155111 | *CDK19* | -1.0 | 9.94E-06 |
| 15 | ENSG00000185862 | *EVI2B* | -1.3 | 1.67E-05 |
| 16 | ENSG00000000971 | *CFH* | -1.0 | 0.0002 |
| 17 | ENSG00000171051 | *FPR1* | -1.3 | 0.0003 |
| 18 | ENSG00000117594 | *HSD11B1* | -1.0 | 0.0008 |
| 19 | ENSG00000189184 | *PCDH18* | -1.2 | 0.001 |
| 20 | ENSG00000134516 | *DOCK2* | -1.0 | 0.002 |
| 21 | ENSG00000007944 | *MYLIP* | -1.2 | 0.006 |
| 22 | ENSG00000092969 | *TGFB2* | -1.2 | 0.007 |
| 23 | ENSG00000187634 | *SAMD11* | -1.2 | 0.02 |
| **Upregulated genes** | | | | |
| 24 | ENSG00000165092 | *ALDH1A1* | 2.0 | 2.32E-37 |
| 25 | ENSG00000197632 | *SERPINB2* | 1.8 | 1.63E-25 |
| 26 | ENSG00000131737 | *KRT34* | 1.5 | 1.00E-23 |
| 27 | ENSG00000154175 | *ABI3BP* | 1.8 | 4.05E-23 |
| 28 | ENSG00000149591 | *TAGLN* | 2.1 | 2.48E-22 |
| 29 | ENSG00000171346 | *KRT15* | 2.9 | 3.80E-20 |
| 30 | ENSG00000164283 | *ESM1* | 1.5 | 1.85E-14 |
| 31 | ENSG00000162892 | *IL24* | 2.0 | 2.57E-12 |
| 32 | ENSG00000164619 | *BMPER* | 1.5 | 6.17E-10 |
| 33 | ENSG00000082482 | *KCNK2* | 1.1 | 7.30E-10 |
| 34 | ENSG00000170961 | *HAS2* | 1.3 | 7.85E-10 |
| 35 | ENSG00000139874 | SSTR1 | 1.1 | 8.26E-10 |
| 36 | ENSG00000123496 | *IL13RA2* | 1.1 | 1.64E-08 |
| 37 | ENSG00000182752 | *PAPPA* | 1.4 | 1.64E-08 |
| 38 | ENSG00000081041 | *CXCL2* | 1.5 | 2.00E-08 |
| 39 | ENSG00000169429 | *IL8* | 1.3 | 6.25E-08 |
| 40 | ENSG00000163734 | *CXCL3* | 1.2 | 8.70E-07 |
| 41 | ENSG00000164220 | *F2RL2* | 1.4 | 1.24E-06 |
| 42 | ENSG00000117152 | *RGS4* | 1.1 | 6.14E-06 |
| 43 | ENSG00000183454 | *GRIN2A* | 1.3 | 3.81E-05 |
| 44 | ENSG00000170989 | *S1PR1* | 1.0 | 4.04E-05 |
| 45 | ENSG00000163739 | *CXCL1* | 1.1 | 4.31E-05 |
| 46 | ENSG00000186847 | *KRT14* | 1.9 | 0.0002 |
| 47 | ENSG00000164161 | *HHIP* | 1.1 | 0.0002 |
| 48 | ENSG00000144821 | *MYH15* | 1.0 | 0.0007 |
| 49 | ENSG00000145358 | *DDIT4L* | 1.1 | 0.001 |
| 50 | ENSG00000152056 | *AP1S3* | 1.1 | 0.003 |
| 51 | ENSG00000196611 | *MMP1* | 1.9 | 0.006 |
| 52 | ENSG00000115008 | *IL1A* | 1.1 | 0.007 |
| 53 | ENSG00000126010 | *GRPR* | 1.1 | 0.01 |
| 54 | ENSG00000163823 | *CCR1* | 1.2 | 0.01 |
| 55 | ENSG00000189223 | *PAX8-AS1* | 1.3 | 9.22E-05 |
| 56 | ENSG00000189001 | *SBSN* | 1 | 4.78E-08 |
