## Supplementary table 6 for "Dissecting *ARL15* Function in Rheumatoid Arthritis: Insights from *Ex Vivo* and *In Vitro* Synovial Fibroblast Models"

| **Supplementary table 6:** Shows expression levels (FPKM) of top ten differentially expressed genes between caucacian (CEU) RASF and CEU healthy SF (as reported in Wang et al., 2014); and corresponding FPKM values in two RASF samples RASF3 (synovial fluid derived) and RASF6 (Synovial tissue derived) analysed in this study. An overall similarity in the trend is observed. |
| --- |

| **Expression levels of top 10 genes in CEU RA & Healthy us RASF3 & RASF6** | | | | |
| --- | --- | --- | --- | --- |
| **Gene** | **CEU RASF** | **CEU_HEALTHY SF** | **RASF3** | **RA6ST** |
| ***CHI3L1*** | **5,847.72** | **0.21** | **258.56** | **85.17** |
| ***MMP1*** | **78.27** | **0.02** | **1.51** | **5.62** |
| ***SMOC2*** | **36.80** | **0.01** | **0.24** | **16.46** |
| ***Ror2*** | **9.66** | **0.01** | **0.05** | **1.67** |
| ***VIT*** | **9.31** | **0.01** | **19.13** | **11.65** |
| ***HEYL*** | **0.04** | **28.46** | **0.35** | **0.01** |
| ***MCAM*** | **0.16** | **153.67** | **0.96** | **2.68** |
| ***EFHD1*** | **0.05** | **103.32** | **1.62** | **0.09** |
| ***FOXE1*** | **0.01** | **18.43** | **0.03** | **0.04** |
| ***WFDC1*** | **0.02** | **76.10** | **0.03** | **0.08** |
