## Supplementary table 7 for "Dissecting *ARL15* Function in Rheumatoid Arthritis: Insights from *Ex Vivo* and *In Vitro* Synovial Fibroblast Models"

| **Expression of genes already reported to be affected upon *ARL15* knock down in RASF (Kashyap et al., 2018)** | | | | | | | | | |
| --- | --- | --- | --- | --- | --- | --- | --- | --- | --- |
| **Number** | **Gene name** | **RASF3 Ctrl FPKM** | **RASF3 ARL KD FPKM** | **log2(fold change) RA3** | **P_value_ (RA3)** | **RASF6 Ctrl FPKM** | **RASF6 ARL KD FPKM** | **log2(fold change) RA6** | **P_value_ (RA6)** |
| **1** | **IL6** | **5.83** | **2.41** | **-1.27** | **0.0243** | **12.12** | **9.63** | **-0.33** | **0.4225** |
| **2** | **ADIPOQ** | **0.02** | **0.00** | **NA** | **NA** | **0.00** | **0.01** | **NA** | **NA** |
| **3** | **ADIPOR1** | **34.05** | **36.24** | **0.09** | **0.7174** | **47.39** | **49.28** | **0.06** | **0.85335** |
| **4** | **PLD1** | **4.68** | **2.18** | **-1.10** | **0.05** | **3.44** | **3.69** | **0.10** | **0.86715** |

**Supplementary table 7: *IL6* was downregulated in both RA samples, adiponectin was upregulated in fluid derived but downregulated in tissue derived RASF. Adiponectin showed similar trend as reported but not significantly and PLD1 was down regulated in fluid derived RASF only.**
