## Supplementary table 8 for "Dissecting *ARL15* Function in Rheumatoid Arthritis: Insights from *Ex Vivo* and *In Vitro* Synovial Fibroblast Models"

**Supplementary Table 8: qPCR primers used for *ARL15* KD confirmation and validation of transcriptome findings (Is this only for MH7A??)**

**Add RASF primers**

| **S.No** | **Gene** | **Forward/**  **Reverse** | **Primer sequence**  **(5’-------3’)** | **Product size (bp)** |
| --- | --- | --- | --- | --- |
| 1 | *ARL15* | Forward | CATCAAGACAAGCCAGCAGC | 113 |
|  |  | Reverse | ATGTCATCCAGTGAGCAGGG |  |
| 3 | *Actin* | Forward | GGCACCCAGCACAATGAAGA | 108 |
|  |  | Reverse | CACATCTGCTGGAAGGTGGA |  |
| 4 | *IL8* | Forward | GCAGAGGGTTGTGGAGAAGTT | 143 |
|  |  | Reverse | AAAGGCAGATACCTAATGACGAT |  |
| 5 | *HAS2* | Forward | ACAGGCATCTCACGAACCG | 120 |
|  |  | Reverse | AACGGGTCTGCTGGTTTAGC |  |
| 6 | *IL24* | Forward | CACCCTGCTGGAGTTCTACTT | 127 |
|  |  | Reverse | TTGCAGTTGTGACACGATGAG |  |
| 7 | *TMED10* | Forward | ATGCGTGATACCAACGAGTCA | 125 |
|  |  | Reverse | TTCTTGGCCTTGAAGAAGCGT |  |
| 8 | *RARRES2* | Forward | CAGGCCCAATGGGAGGAAAC | 103 |
|  |  | Reverse | AACTTGGGTCTCTATGGGGC |  |
| 9 | *UBC* | Forward | ATTTGGGTCGCGGTTCTTG | 133 |
|  |  | Reverse | TGCCTTGACATTCTCGATGGT |  |
| 10 | *ESM1* | Forward | GGTATCTGCAAAGAGCATGACA | 100 |
|  |  | Reverse | ATTTCCTCATTACGGGAGACCC |  |
| 11 | *TFPI* | Forward | GAATAACTCCCTGACTCCGCAA | 116 |
|  |  | Reverse | ATCTGTTCTCATTGGCACGACA |  |
| 12 | *PAPPA* | Forward | GTCTCAAGTGGTATCCTCACCC | 111 |
|  |  | Reverse | CATAGTTGCAAAAGGCTCGGTT |  |
| 13 | *IL1A* | Forward | CACAGGTAGTGAGACCAACCTC | 114 |
|  |  | Reverse | ACCCAGTAGTCTTGCTTTGTGG |  |
| 13 | *MSN* | Forward | AGAGCCTGCTGAGAATGAGC | 108 |
|  |  | Reverse | GTACGTTCCTCCTCACTGCG |  |
| 14 | *FPR1* | Forward | TGCTCCTCACATTGCCAGTTAT | 109 |
|  |  | Reverse | CTCTCTTTAGGGTCGTTGGTCC |  |
| 15 | *DOCK2* | Forward | GGAATTAGCATCACCCAAGACG | 119 |
|  |  | Reverse | CTACGCTGCTTCTCTTTGTCCT |  |
| 16 | *TGFB2* | Forward | AGGATAATTGCTGCCTACGTCC | 100 |
|  |  | Reverse | GCACAGAAGTTGGCATTGTACC |  |
| 17 | *CD82* | Forward | GCTCATTCGAGACTACAACAGC | 108 |
|  |  | Reverse | GTCCAGTTGTAGAAGCTGACCC |  |
| 18 | *TLR4* | Forward | TTCAGCTCTGCCTTCACTACAG | 101 |
|  |  | Reverse | CACAACAATCACCTTTCGGCTT |  |
| 19 | *MMP1* | Forward | GGACAGAATGTGCTACACGGATA | 100 |
|  |  | Reverse | TGTTTTCCTCAGAAAGAGCAGCA |  |
| 20 | *S1PR1* | Forward | CTCGGTCTCTGACTACGTCAAC | 108 |
|  |  | Reverse | ACCGAGGTCAGTTTAATGCTGT |  |
| 21 | *GRPR* | Forward | CACAGCACTGGAAGGAGTACAA | 105 |
|  |  | Reverse | ATACCGCTCGTGACAGATGTTT |  |
| 22 | *CCR1* | Forward | ATGACACGACCACAGAGTTTGA | 105 |
|  |  | Reverse | ATACCAAGGAGTACAGAGGGGG |  |
| 23 | *CXCL2* | Forward | AAACCGAAGTCATAGCCACACT | 110 |
|  |  | Reverse | GTTGGATTTGCCATTTTTCAGCA |  |
| 24 | *CXCL3* | Forward | CACACTCAAGAATGGGAAGAAAGC | 96 |
|  |  | Reverse | CAGTTGGTGCTCCCCTTGTT |  |
